# Hippocampal conditioning code dominates and disrupts the place code

**DOI:** 10.64898/2026.02.17.706430

**Authors:** Hannah S Wirtshafter, Mayank R Mehta, Sara A Solla, John F Disterhoft

**Affiliations:** Department of Neuroscience, Northwestern University Feinberg School of Medicine, Chicago, IL 60611, USA; W. M. Keck Center for Neurophysics, Departments of: Physics & Astronomy, Neurobiology, Neurology, University of California, Los Angeles, Los Angeles, CA 90095, USA; Department of Physics and Astronomy, Northwestern University, Evanston, IL 60208, USA

## Abstract

The hippocampus is widely viewed as a spatial map because many CA1 neurons exhibit spatially selective activity during exploration. However, the hippocampus is also involved in nonspatial learning, including conditioning tasks. Most prior reports of nonspatial hippocampal activity were obtained in immobile animals, leading to the proposal that the hippocampus encodes spatial variables during locomotion and nonspatial variables during immobility. The question of how the hippocampus encodes both spatial and nonspatial variables during locomotion and spatial exploration has remained unanswered. To address this issue directly, we used calcium imaging to record the activity of thousands of CA1 neurons from rats tested on a conditioning task while freely exploring an open field. Across more than 6,000 neurons from five rats, the rate of calcium events increased during conditioning task periods relative to non-task periods, an effect that persisted after controlling for locomotor speed. At the single cell level, conditioning task modulation was widespread and strongly biased toward increased activity; neurons whose activity was enhanced during the conditioning task outnumbered spatially selective neurons by more than threefold. In addition, many neurons showed significantly increased activity during specific segments of the conditioning task. At the population level, conditioning coding remained invariant to changing spatial location; in contrast, spatial coding was reduced during task periods. These findings show that during active behavior, engagement in the conditioning task reorganizes hippocampal activity into a temporally structured pattern of population activity that can dominate and disrupt place coding. Together, these results challenge the view that the hippocampus functions primarily as a stable spatial map.

## Introduction

Since the discovery of place cells over 50 years ago^1–3^, the hippocampus is widely regarded as a spatial map because many CA1 neurons exhibit location-specific activity during exploration^4–9^. However, the hippocampus is also required for nonspatial cognitive tasks^10–14^ and exhibits task-related activity during these behaviors^15–29^, including trace eyeblink conditioning (tEBC), a hippocampus-dependent task commonly used to study associative memory^16,17,19,30–35^. Because most recordings of nonspatial hippocampal activity were obtained in immobile or head-fixed animals^15,16,36,37^, it has been proposed that spatial variables dominate hippocampal activity during locomotion, whereas nonspatial variables are represented primarily during immobility^5,6,38,39^. Many studies of hippocampal involvement in nonspatial tasks tightly couple task structure to spatial position^31,40^, limiting the ability to dissociate the encoding of spatial and nonspatial variables. Few studies have addressed this question, with mixed and sometimes conflicting results^13,38^. Whether nonspatial activity during active exploration reflects genuine task encoding rather than indirect modulation of spatially related activity remains unresolved.

To address this question, we recorded large populations of CA1 neurons using calcium imaging while rats performed trace eyeblink conditioning (tEBC) during free exploration of an open field. This experimental design allowed us to dissociate spatial position, movement, and conditioning task structure while monitoring hippocampal activity at both the single cell and population levels.

## Results

We trained a cohort of five freely moving rats on the tEBC task while recording neural activity from the dorsal CA1 region using miniature microscopes and the calcium indicator GCaMP8m. In this task, each tEBC trial consists of a 250ms auditory tone (conditioned stimulus, CS), followed by a 500ms trace interval with no stimulus, and a 100ms eyelid shock (unconditioned stimulus, US) (Fig. 1a-b). Intertrial intervals were randomly varied between 30 and 60 seconds (mean 45 seconds). Blink responses and shock delivery were monitored using EMG wires implanted in the eyelid muscle (see Methods). Throughout training and testing, animals were allowed to move freely within the behavioral arena (78.7cm x 50.8cm box); animals did not exhibit freezing or pausing during trials (Fig. 1c). Data were analyzed only in those sessions where the rats met the behavioral criterion for task acquisition: they performed tEBC on at least 70% of trials across three consecutive sessions (see Methods, Fig. S1).

**Figure 1.**
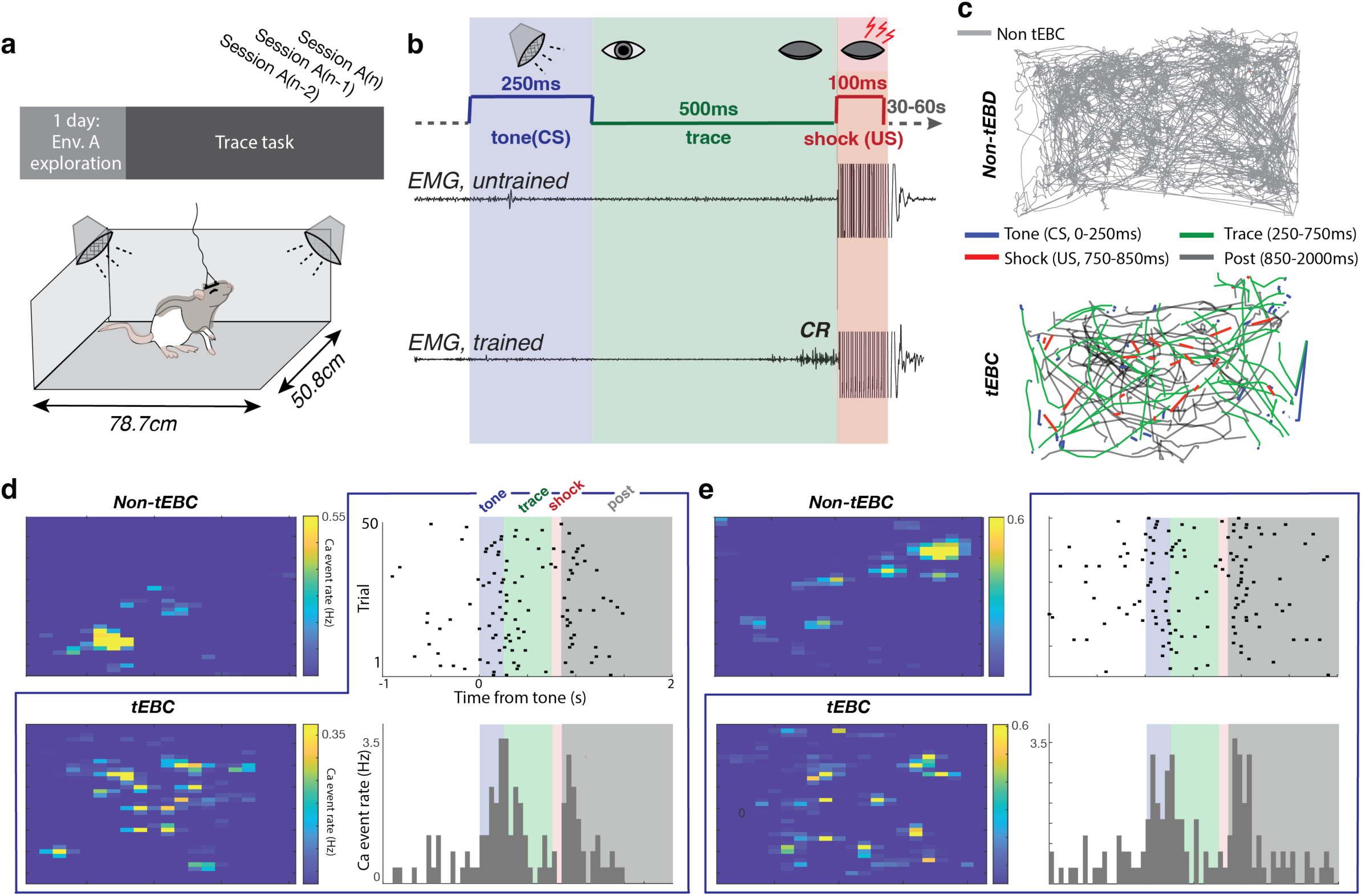
Schematic of task and cell responses. **a.** Schematic of the trace eyeblink conditioning (tEBC) environment. After one day of exploration in the conditioning environment, rats were trained daily in tEBC until reaching the behavioral criterion for task acquisition (see Methods). **b.** Schematic of the trace eyeblink conditioning task. Trial timeline: a 250ms tone (auditory conditioned stimulus, CS), a 500ms trace interval, and a 100ms eyelid shock (unconditioned stimulus, US). EMG recordings from the orbicularis oculi muscle illustrate untrained (top) and trained (bottom) animals; the shock period saturates the EMG trace, truncated here for visualization. Conditioned responses (CRs) are observed in trained animals prior to US onset. **c.** Representative movement trajectories during conditioning sessions. Top: Gray traces depict trajectories during times when the animal is not performing the tEBC task. Bottom: Blue trajectories depict movement during the tone (CS), green during the trace, red during the shock (US), and black in the post-shock period of the tEBC task. **d.** Example neuron showing rate maps in the environment both during no tEBC task (top left) and tEBC task (bottom left) periods. Activity during tEBC periods is also represented as raster plots (top right) and as peristimulus time histograms (PSTH, bottom right) aligned to tone (CS) onset. Boxed graphs indicate figures corresponding to the tECB task periods. **e.** Same as d but for a different neuron.

We recorded and analyzed a total of 6192 dorsal CA1 cells from five rats across 15 sessions, three sessions per rat (See Methods). Many neurons showed clear spatial modulation, consistent with extensive literature on place cells^2,41,42^ (Fig. 1d,e, upper-left panels, S2). Simultaneously recorded neurons showed distinct responses during different phases of the task, with their activity aligned to the tone, trace interval, eyelid shock, or post-shock period; these responses were stable across trials (Fig. 1d-e, S2). During tEBC, the rats occupied diverse locations in the arena, yet the neurons were active selectively and reliably during specific epochs of the trace task, showing that the trace responses were not merely spatial responses. Neural activity during the tEBC task was so reliable that the neuron’s spatial selectivity was substantially degraded and differed compared to the non-tEBC periods. We performed several analyses and controls to confirm these observations, as follows.

### Hippocampal neurons encode distinct aspects of the tEBC task

To assess how hippocampal activity during the tEBC task differs from ongoing activity during free exploration, we quantified calcium event rates separately during tEBC and non-tEBC periods. At the population level, calcium event rates during the tEBC task were positively correlated with non-task rates (Fig. 2a), indicating that baseline firing tendencies were preserved across behavioral contexts. However, activity during task periods was systematically elevated for most cells. This increase was not driven by a small subset of highly active neurons but reflected a general shift across the population (Fig. 2a; Fig. S3–4).

**Fig 2.**
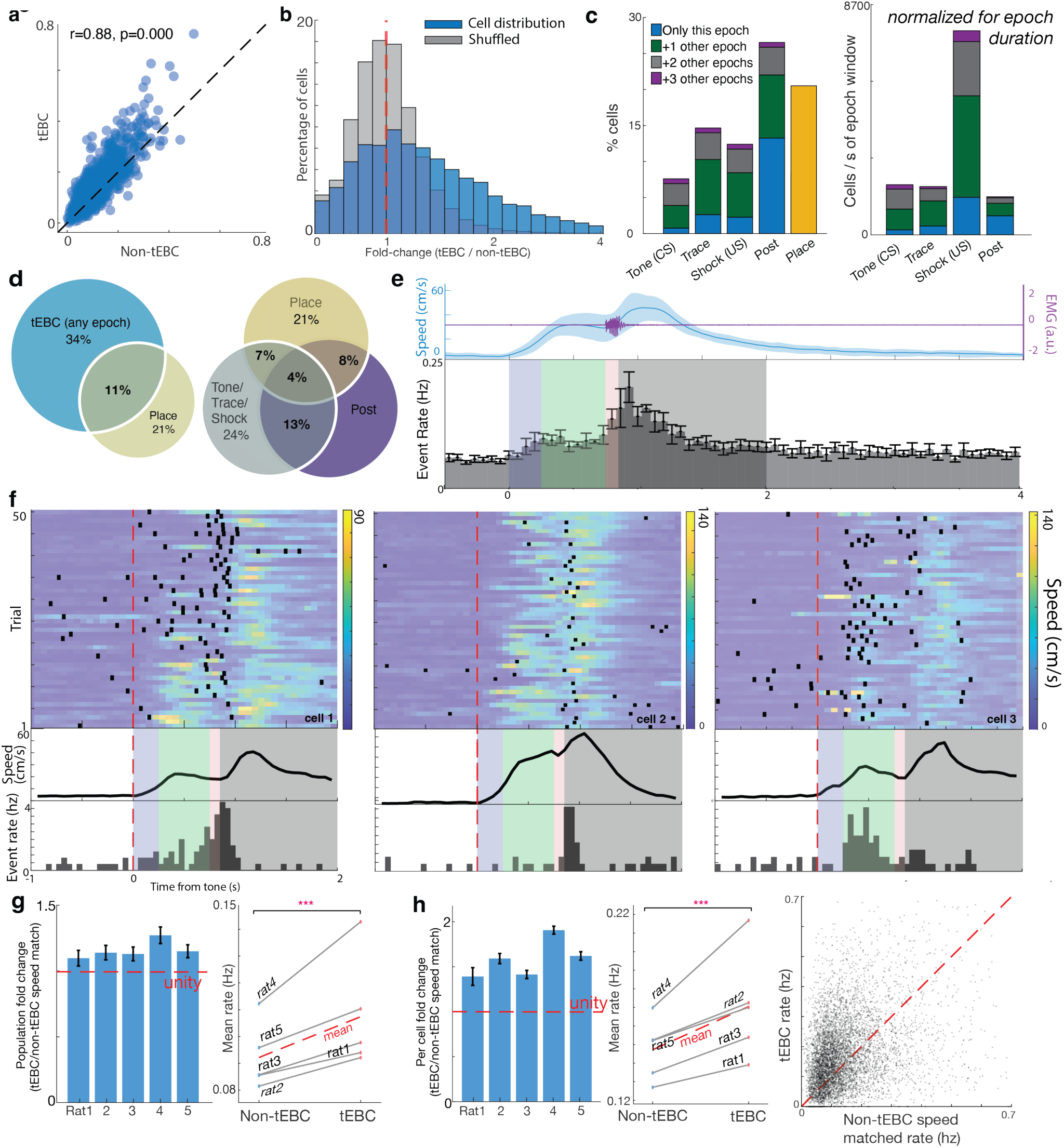
**a.** Positive correlation between calcium event rates during tEBC and non-tEBC periods, pooled across all animals. The majority of points fall above the dash diagonal identity line, indicating an overall higher rate during tEBC periods. Pearson’s r and p-value as shown. See also figure S3 and S4. **b.** Distribution of event rate change (tECB periods /non-tEBC periods) across all recorded cells (blue) compared to a shuffled null distribution (gray). The dotted red line indicates no change (fold-change = 1). Modulation was biased toward increases, with 34.2 ± 8.2% of cells exhibiting significant task-enhanced activity (one-sided permutation test for increases, α = 0.05). Plot ranges are set to the 99.5th percentile of pooled observed and shuffled values for visualization. **c.** Proportion of cells significantly modulated by tEBC; stacks show how many also have substantial modulation in +1/+2/+3 other epochs. Left: For each task epoch (Tone/CS, Trace, Shock/US, Post) and for place, bars show the percent of cells significantly modulated. The count data has been normalized so that each rat contributes equally. Stacks partition each epoch’s modulated population into cells modulated only in that epoch (blue) or additionally in **+**1 (green), +2 (gray), or +3 (purple) other epochs. Right: Bars show again the percent of cells significantly modulated in each epoch. The cell count data has been normalized by epoch duration (cells per second of the epoch window) to compensate for large differences in window length. Taller bars therefore indicate higher rate of modulation, independent of how long the epoch lasts. See Fig. S5 for the same analysis restricted to place cells only. Epoch membership proportions were broadly similar to those observed in the full population, although a higher proportion of place cells showed post modulation (paired t-tests across rats, Bonferroni corrected, post p < 0.05). **d.** Venn diagrams showing overlap of tEBC selectivity and spatial selectivity. The data has been normalized so that each rat contributes equally. Left: Venn diagram shows percentage of cells modulated by tEBC versus place. Right: Cell selectivity when ‘tone+trace+shock’ are treated as one epoch and ‘post-shock’ as a separate epoch. Percentages indicate the proportion of cells in each category; circle areas are not to scale. **e.** Top: Trial-aligned averages of running speed (blue, left y axis) and EMG activity (purple, right y axis) across all sessions and all rats. Purple shaded EMG activity spans ±SEM. Rats maintained locomotion throughout the tEBC period; EMG responses reliably occurred during US delivery. Bottom: peristimulus time histogram (PSTH) averaged across all rats, showing the average calcium event dynamics of CA1 neurons during the tEBC task. The mean calcium event rate is shown for successive 133ms bins from 1s before the tone (CS) onset to 2s after the end of the post-shock epoch; error bars denote the SEM across cells. See figure S6 for individual animals and figure S7 for comparison of task vs pre task running speed. **f.** Top row: Trial-by-trial raster plots (black dots) overlaid on trial-aligned running speed heatmaps for three example neurons. Red dashed line indicates tone (CS) onset. Middle row: Average speed across all trials for the corresponding session for that rat. Bottom row: PSTH for the three shown cells. Calcium events occurred across a wide range of speeds and cells were tuned to different epochs within the task. **g.** Comparison of population-level calcium event rates during tEBC periods versus speed-matched non-tEBC periods. Population rates were computed by pooling calcium events across all neurons within a session and dividing by total time. Left: Fold change in calcium event rate (tEBC /non-tEBC, speed-matched) for each rat, shown as mean ± SEM for the neural population. The red line marks the across-animal mean. At the population level, there were on average 1.16x more calcium events during tEBC periods than during speed-matched non-tEBC periods (individual rats mean ± SEM: 1.10 ± 0.04 1.14 ± 0.05 1.13 ± 0.05, 1.27 ± 0.06, and 1.15± 0.05). Right: For each rat, mean calcium event rates during non-tEBC and tEBC periods. Gray lines connect matched values from the same rat; the red line indicates the across-animal mean. Across rats, calcium event rates were significantly higher during the tECB periods than during speed-matched non-tEBC periods (paired t-test, ***p<0.001). See figure S9 for an analogous analysis based on acceleration data. **h.** Comparison of calcium event rates at the level of individual neurons during tEBC periods versus speed-matched non-tEBC periods. Calcium event rates were computed separately for each neuron, and statistics were then performed across the population of neurons. Left: Fold change in firing rate (tEBC /non-tEBC) for each rat, shown as mean ± SEM across tested cells. The red line marks the across-animal mean. Neurons had on average 1.68x more calcium events during tEBC periods than during speed-matched non-tEBC periods (individual mean ± SEM:1.39 ± 0.10, 1.59± 0.05, 1.41 ± 0.04, 1.91 ± 0.05, and 1.62 ± 0.05. Middle: For each rat, mean firing rates during tEBC and non-tEBC periods. Gray lines connect matched values from the same rat; the red line indicates the across-animal mean. Across rats, calcium event rates were significantly higher during the tEBC periods than during speed matched non-tEBC periods (paired t-test, ***p<0.001). Right: scatter plot comparing calcium event rates during tEBC period to the rates during speed-matched non-tEBC periods within a session. The majority of points fall above the dash diagonal identity line, indicating an overall higher rate during tEBC periods. See figure S10 for an analogous analysis based on acceleration data.

We next asked how many neurons exhibited significant increases in activity during tEBC periods. Comparing calcium event rates during the entire tEBC task window to intertrial periods of the same duration revealed that 34.2 ± 8.2% of CA1 neurons exhibited significant task-enhanced activity relative to a shuffled null distribution (one-sided permutation test, α = 0.05; Fig. 2b–c). A smaller fraction (4.2% ± 2.8%) showed significant task suppression. This directional bias was evident at the population level, consistent with a net recruitment of hippocampal cells during conditioning.

We then examined the temporal profile of task-modulated neurons. Among cells that were significantly modulated across the full tEBC window, responses were not uniformly distributed across the task period. Most neurons showed peak modulation in a specific epoch (CS, trace, US, or post-shock), indicating temporally specific encoding rather than diffuse task engagement (Fig. 2c, Fig. S5). This result demonstrates that neurons encoded specific epochs of the tEBC task rather than nonspecific mechanisms such as altered arousal state.

Task-modulated neurons exhibited pronounced temporal heterogeneity. Distinct subsets of cells were preferentially active during the tone, trace, shock, or post-shock epochs, with relatively limited overlap across epochs (Fig. 2c, f, Fig. S5–S10). We found that 85.7% of significantly task-modulated neurons were selective for one or two epochs, rather than being broadly active throughout the full task period, indicating that the tEBC task is neuronally structured in time. The post-shock period contained the largest absolute fraction of modulated cells. Note that this epoch was also substantially longer than the others—a short 100ms shock epoch versus the longer 1.15s post-shock epoch. After normalizing modulation by epoch duration, the density of post-shock modulation was comparable to that observed during the tone and trace epochs; all three were exceeded by modulation during the brief shock epoch (Fig. 2c). The apparent dominance of post-shock responses reflects window length rather than preferential recruitment.

Overlap analyses showed that modulated activity during tEBC periods was not restricted to a distinct cell class (Fig. 2d). Task-selective and place-selective populations partially overlapped, consistent with the idea that the trace task recruits neurons that may also participate in spatial coding. Spatial tuning is examined in detail later (Fig. 4); the overlap presented here is only interpreted as indicating that task modulation is not confined to non-place cells.

An increase in running speed during the trace period at the population level (Fig. 2e, Fig. S6–11) raises the possibility that elevated neural activity during this period reflects a global change in behavioral state. If this possibility held, it would lead to two predictions: first, if tEBC-related activity reflected speed modulation, then neural responses should scale with speed both within and across trials, and second, differences between task and non-task periods should disappear when controlling for speed. The analyses presented in the remaining panels of Figure 2 show that neither of these predictions holds.

Examples of single cell activity across trials revealed that tEBC responses were heterogeneous and temporally structured, rather than reflecting a uniform gain change (Fig. 2f). Individual neurons exhibited diverse response profiles, including transient responses aligned to tone or shock, ramping activity during the trace interval, and delayed or prolonged post-shock responses. Importantly, these calcium events occurred across a wide range of running speeds, indicating that the activity changes of individual cells are not trivially explained by a correlation with running speed. Additionally, correlations between running speed and calcium event rate were near zero across cells, both at the single-trial level and within trials, and were indistinguishable from correlations observed in shuffled controls (Fig. S8). The population-level correlation between running speed and neural activity (Fig. 2e) does not correspond to velocity tuning at the level of individual neurons.

To directly test whether running speed could account for neural activity during tEBC periods, we compared calcium event rate during tEBC periods to those during non-tEBC periods matched for running speed. Even under this stringent control, calcium event rates remained significantly elevated during the task at both the population level (Fig. 2g) and the single cell level (Fig. 2h). Note that the effect was substantially larger at the single cell level; most neurons showed increased calcium event rates during tEBC periods compared to speed-matched non-tEBC periods (mean 1.68x more calcium events during tEBC). These effects were consistent across animals and remained significant relative to permutation controls (p=0, data not shown); similar results were observed after controlling for acceleration (Fig. S9). Together, these findings show that hippocampal neurons exhibit robust, heterogeneous, tEBC-related increases in activity that are not explained by concurrent increases in running speed. The tEBC task engages a broad subset of CA1 cells whose activity increases across multiple tEBC epochs, with distinct temporal profiles spanning the tone, trace, shock, and post-shock epochs. Notably, these increased responses persisted when comparing speed-matched tEBC and non-tEBC periods.

We conclude that the increased responses during tEBC periods are not attributable to locomotor speed. The tEBC task recruits a hippocampal population-level response in which structured firing is not fully accounted for by motor variables.

### tEBC evokes a temporally structured, trajectory-like population code

We next asked how the task organizes population activity. To this end, we used population vectors (PVs) defined in a neural space with one axis for each of the neurons recorded in a given session; each component of the PV quantifies the calcium event rate for the corresponding neuron. A comparison of PVs across behavioral epochs showed that activity during the tEBC period was highly consistent across trials and clearly distinct from non-tEBC activity (Fig. 3a). Population similarity, measured as the Pearson correlation across neurons, was substantially higher within the tEBC period than the similarity between tEBC and non-tEBC periods or within non-tEBC periods. This distinction remained when non-tEBC periods were selected to match the running speed profile of the tEBC periods, indicating that differences in locomotor state alone do not explain the distinct pattern of population vectors observed during the tEBC task. Notably, activity during speed-matched non-tEBC periods was itself consistent; this similarity did not differ significantly from that across tEBC periods, suggesting that both tEBC and non-tEBC states can support stable ensemble structure. However, the tEBC periods were associated with a population configuration that was systematically different from that during non-tEBC periods, reflecting a reproducible, trial-associated ensemble pattern of activity.

**Figure 3.**
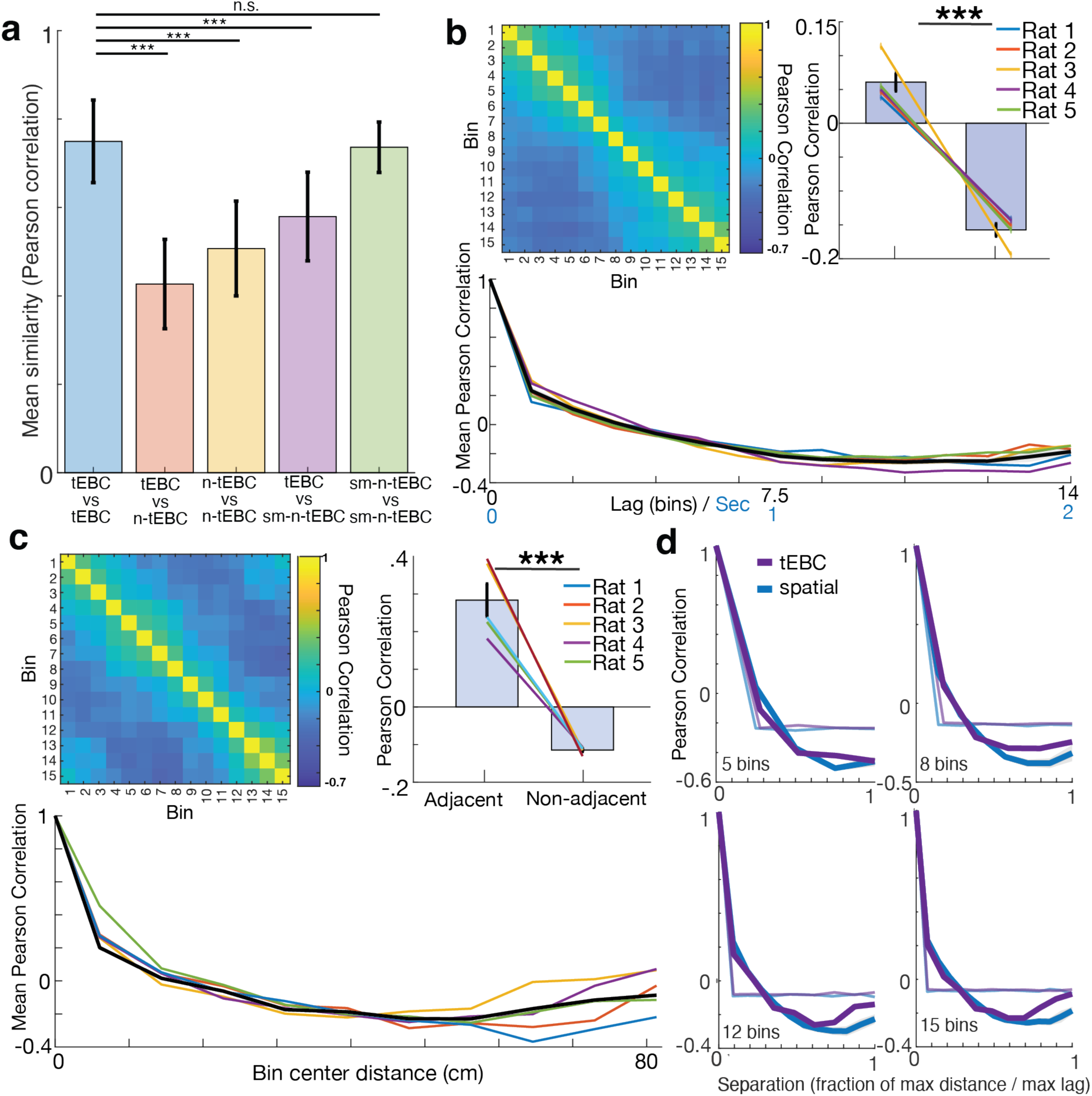
Heterogeneous but distinct neuronal dynamics and coding across trials. **a.** n-tEBC = non-tEBC, sm-n-tEBC = speed-matched non-tEBC task. Bars show session means ± SD. For each session for a given rat, each 2s window associated with a tEBC trial was compared to (i) a 2s windows associated with other tEBC trials, (ii) randomly selected 2s windows involving no tEBC, and (iii) a speed-matched 2s windows drawn strictly from outside tEBC windows. For each bootstrap draw, trials were split into disjoint groups to form population vectors (PVs), and similarity was quantified as the Pearson correlation across neurons. Similarity across tEBC periods (tEBC vs tEBC; 0.750±0.093) was substantially higher than similarity between tEBS and non-tEBC periods (tEBC vs n-tEBC; 0.427±0.101), non-tEBC to non-tEBC periods (n-tEBC vs n-tEBC; 0.507±0.107), and similarity between tEBC periods and speed-matched non-tEBC periods (tECB vs sm-n-tEBC; 0.580±0.100) (paired tests across sessions, ***p < 0.0001 for all comparisons). Note that the activity during speed-matched non-tEBC periods was highly similar (sm-n-tEBC vs sm-n-tEBC; 0.736±0.057). and did not differ significantly from the similarity across tEBC periods. **b.** Population vector similarity during tEBC is time-structured and shared across rats. Top left: similarity matrix for PVs corresponding to different times along a tEBC trial. The 2s duration of a tEBC following the CS onset was divided into equal 15 bins. PVs were built from split-averaged event rates for each neuron and compared using Pearson correlations. The matrix shows a heat map of Person correlations between bins, averaged across rats. A bright diagonal and banded, increasingly dimmer off-diagonals indicate that population activity is most similar for nearby time bins and progressively less similar with temporal distance. Top right: within vs between epochs. Bins were grouped into CS, trace, US, and post-US epochs. Bars show across-rat mean ± SEM similarity for all within-epoch pairs vs between-epoch pairs; colored lines correspond to individual rats. Within-epoch similarity exceeded between-epoch similarity (Paired t test, p*** = <0.001 across all rats and for each rat, see figure S11); this indicates that activity is not only temporally smooth but also clustered by task epoch. Bottom: Lag curves. Mean similarity is plotted as a function of lag (difference in bin index). Colored lines correspond to individual rats; thick black line shows the mean across rats. All rats show a monotonic decay of similarity with lag, consistent with a temporally organized population code. (Fig. S11). Split-half analyses confirmed that lag structures were reliable within session (Fig. S12) **c.** Spatial population similarity is structured by physical proximity. Top left, similarity matrix for PVs corresponding to different positions in the environment. Population vectors (PVs) were built from 15 equal-occupancy spatial bins computed from non-tEBC running epochs at speeds ≥ 4 cm/s. In each bin, PVs were built from mean event rates for each cell and compared across bins using Pearson correlations. The matrix shows a heat map of Person correlations between bins, averaged across rats. The bright diagonal and banded, increasingly dimmer off-diagonals indicate that population activity is most similar for nearby bins that share borders. Top right: spatially adjacent bins vs non-adjacent bins (adjacency defined by cells sharing a side, not along diagonals). Bars show across-rat mean ± SEM similarity; colored lines correspond to individual rats. Adjacent bins were more similar than non-adjacent bins (paired t test, p*** = <0.001 across all rats and for each rat, see figure S13). Bottom: Lag curves. Mean similarity is plotted as a function of lag (distance between bin centers in cm). Colored lines correspond to individual rats; thick black line shows the mean across rats. Similarity generally decreases with physical distance, consistent with a spatial code structured around distance. Split-half analyses confirmed lag structures were reliable within session (Fig. S14). **d.** Temporal and spatial similarity across population vectors show a shared decay profile with distance. Population-vector (PV) similarity measured by the Pearson correlation is plotted versus temporal lag during the tEBC period (0s–2s; temporal, dark purple; one temporal lag bin corresponds to 0.13s) and versus spatial separation during non-trial running (spatial, dark blue; because spatial position is 2D, spatial lag is expressed as distance between the centers of spatial bins rather than the number of bin steps). Each panel shows a different discretization of time/space (5, 8, 12, or 15 bins; top-left to bottom-right). Solid purple and blue lines show means across rats. Faint purple and blue lines provide null references for comparison; these were obtained by shuffling temporal and spatial curves, respectively (mean of 500 permutations in the bin order of the corresponding PV similarity matrix). For every bin resolution, the temporal and spatial curves decline with lag and closely overlap in shape (Pearson r on curves: 0.990, 0.981, 0.975, 0.967 for 5/8/12/15 bins; RMSE_z = 0.123–0.246). The integrated magnitudes of the temporal and spatial curves did not differ from each other (ΔAUC permutation test, p = 0.866, 0.732, 0.624, 0.714 for 5/8/12/15 bins).

To examine the organization within a tEBC trial, we divided the 2s window into 15 identical temporal bins, each of size Δ = 133ms, and computed similarity between PVs from different bins. The resulting similarity matrix, averaged across rats, showed a strong diagonal with banded off-diagonals; lag–similarity curves showed monotonic decay with increasing temporal separation between bins for every rat (Fig. 3b, S11-S12). Population activity was also segmented by task epoch; within-epoch similarities exceeded between-epoch similarities across all rats (Fig. 3b, S11-S12). These results indicate that each tEBC trial engages a conserved, temporally organized trajectory through the neural space.

We next asked whether this temporal trajectory mirrors the distance-dependent structure of spatial coding. We partitioned the arena into spatial bins based on equal occupation over time and computed the similarity between PVs associated with different spatial bins. Spatial PV similarity was highest for adjacent locations and declined with spatial distance, consistent with place coding (Fig. 3c, S13-S14). Strikingly, temporal and spatial lag curves exhibited nearly identical shapes across a range of bin numbers (4-14 bins; Fig. 3d), and split-half analyses confirmed the reliability of both structures (S12, S14). This alignment shows that population vector similarity falls off with “distance” along the relevant axis—time during tEBC, space during exploration.

Together, these results reveal that tEBC engages diverse temporal coding strategies, including early CS-driven responses, trace-interval modulations, US rate increases, and sustained post-US changes. Despite this heterogeneity, population activity during the tEBC period is highly stereotyped and distinct from non-tEBC periods, indicating the emergence of a structured ensemble code during the tEBC task. These results show that trace conditioning restructures CA1 by recruiting neural activity into a robust, temporally organized population code that is distinct from place coding. These two codes, though distinct, involve CA1 population states that obey a similar decay rule: in time during tEBC and in space during exploration. This similarity suggests a shared organizing principle for spatial and temporal hippocampal codes.

### tEBC disrupts hippocampal spatial coding

Neurons exhibited spatially structured calcium activity during free exploration, as quantified by spatial mutual information (MI) computed outside tEBC periods (Fig. S16), consistent with prior reports^43,44^. However, including task-related activity substantially altered this spatial structure (Fig. 4a). When MI was computed using all calcium events, spatial information was reduced relative to estimates obtained after excluding tEBC periods (Fig 4a). Removing calcium events during tEBC periods increased spatial MI across all rats, whereas removing an equivalent number of events from non-tEBC periods, either randomly or matched for running speed, produced little or no increase. These results indicate that activity during tEBC preferentially degrades spatial information, consistent with task-related activity reflecting non-spatial structure.

**Figure 4.**
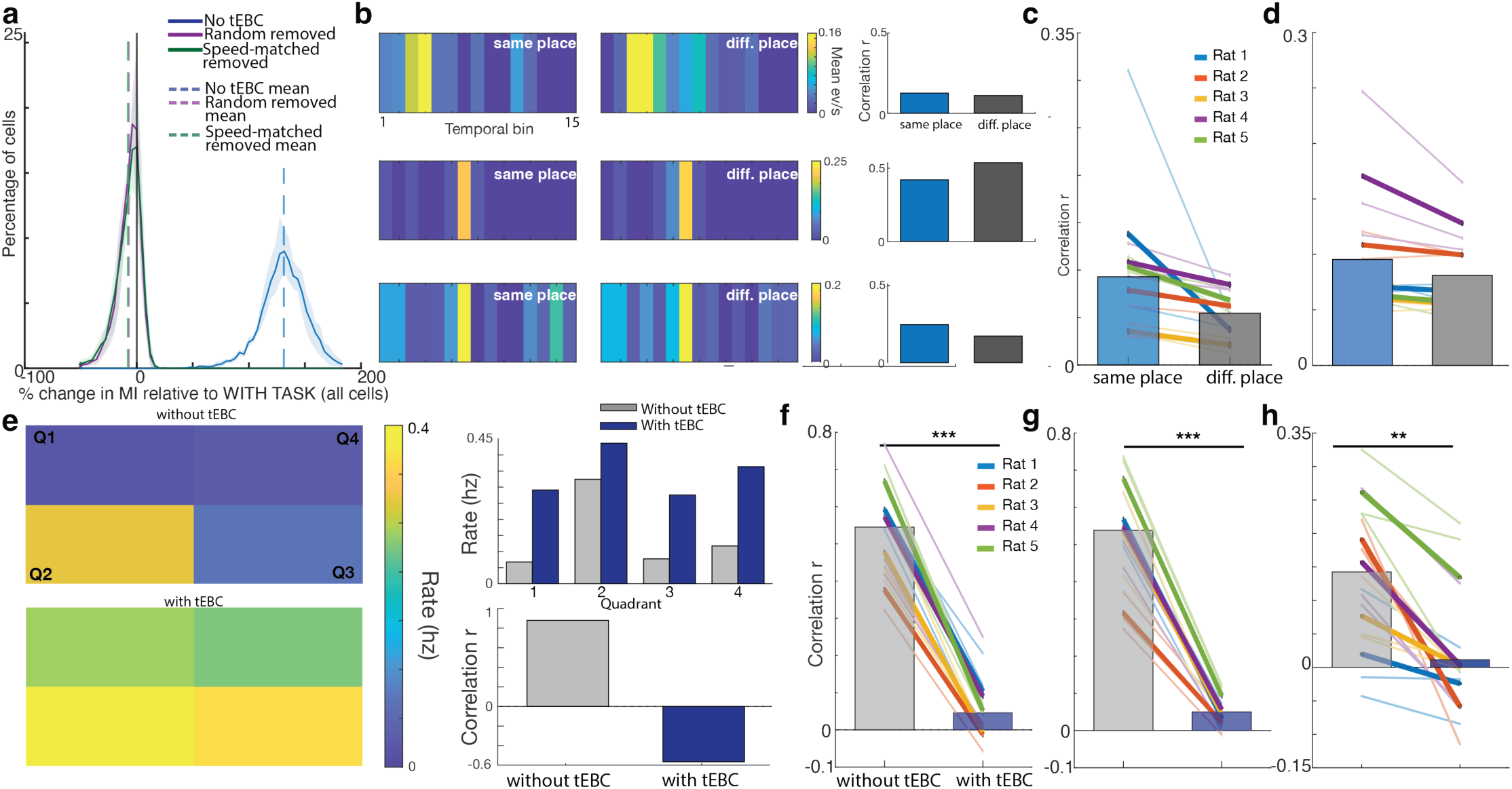
a. Spatial mutual information (MI) compared across four conditions: (1) with tEBC (all calcium events included, baseline), (1) without tEBC (task-period events removed, blue), (3) random removal (an equal number of events removed from non-tEBC periods, purple), and (4) speed-matched removal (an equal number of events removed from non-tEBC periods matched for running speed, green). The distributions show the percent change in MI relative to baseline (with tEBC), computed per cell and averaged equally across animals (shaded regions, 95% bootstrap confidence intervals). The removal of events occurring during tEBC periods resulted in a large increase in spatial MI (130% to 150% increase relative to baseline across rats). In contrast, the removal of an equivalent number of events from non-tEBC periods, either randomly selected or matched for speed, produced minimal changes in MI (0% to 10%). Differences between including tEBC vs excluding tEBC periods were highly significant across cells (paired t-tests, all rats p ≈ 0; pooled t(6191) = −184.3), whereas differences between including tEBC and control removals were small despite statistical significance. See figure S17 for individual rats. b. Single cell temporal activity profiles underlying the spatial similarity analyses in figure 4b-c. Each row shows one example hippocampal neuron from a representative recording session. Left and middle: Mean calcium event rate across the 15 temporal bins within the tEBC period, computed separately for each spatial quadrant by iteratively comparing trials in a given quadrant (e.g., Q1) to the average across trials in the other three quadrants (Q2–Q4). Right: Correlation of temporal activity profiles (Pearson r) for same quadrant to different quadrant comparisons, averaged across quadrants. Note that trial-to-trial consistency of temporally structured activity during tEBC can differ depending on spatial context; this observation motivates the population level split-half similarity analyses shown in panels e–f. c. Similarity of population vectors (PV) for pairs of tEBC trials occurring in the same versus different spatial quadrants. Thin colored lines show session means for each rat (n = 3 sessions per rat), thick colored lines show the mean across sessions for each rat, and bars show the mean across rats (n = 5). PV similarity was only weakly higher for same-quadrant compared to different-quadrant trials. This difference was small and not significant when assessed across rats (paired t test in Fisher z space, t(4) = 2.37, p>0.05), and only modestly significant when assessed across sessions (paired t test in Fisher z space: t(14)=2.30, p=0.04). d. Similarity of single-cell activity for pairs of tEBC trials occurring in the same versus different spatial quadrants. For each cell, similarity was quantified by splitting trials into two halves and computing the correlation of temporal activity profiles across halves. Thin lines, thick lines, and bars are as in panel b. Same-quadrant similarity was only modestly higher than different-quadrant similarity; this effect was small and did not reach significance when assessed across rats (paired t test in Fisher z space, t(4)=1.96, p>0.05), but was modestly significant across sessions (paired t test in Fisher z space, t(14)=2.44, p<0.05). e. Spatial structure of activity for an example hippocampal neuron. Left: Mean calcium event rate across spatial quadrants during non-tEBC (top) and tEBC (bottom) periods. During non-tEBC periods, activity is concentrated in a single quadrant, whereas during tEBC periods it is more distributed across quadrants. Right: Quantification of spatial structure for this neuron. Bars show the similarity of activity patterns across quadrants, computed as the Pearson correlation between quadrant-specific activity vectors. Spatial similarity is high during non-tEBC periods and reduced during tEBC, reflecting a loss of location-specific activity. f. Spatial similarity of population vectors within the same quadrant during non-tEBC and tEBC periods. Thin lines, thick lines, and bars are as in panel b. Spatial similarity was markedly reduced during tEBC compared to non-tEBC periods across sessions and animals (mean r: 0.050 tEBC vs 0.54 non-tEBC). This reduction was robust across animals (paired t-test in z space: t(4)=−10.10, p<0.00) and across sessions (paired t-test in z space: t(14)=−13.81, p<0.001). Significance was further confirmed using permutation-based tests (see Methods). g. Reliability of single-cell spatial coding relative to a baseline spatial map. Spatial activity was quantified across quadrants; for each cell, similarity was computed between its activity map during tEBC and a reference map derived from non-tEBC periods. Thin lines, thick lines, and bars are as in panel b. Single-cell spatial similarity relative to the baseline map was markedly reduced during tEBC (mean r: 0.050 tEBC vs 0.530 non-tEBC). This reduction was robust across animals (paired t-test in z space: t(4)=−7.96, p***=0.001) and across sessions (paired t-test in z space: t(14)=−12.07, p<0.001). Significance was further confirmed using permutation-based tests (see Methods). h. Spatial stability of single cells. Spatial stability was measured as the correlation of each cell’s spatial activity map across quadrants within the same condition (non-tEBC or tEBC). Thin lines, thick lines, and bars are as in panel b. Spatial stability was significantly reduced during tEBC compared to non-tEBC periods. Single cell spatial similarity was significantly lower during tEBC compared to non-tEBC periods (mean r: 0.314 tEBC vs 0.557 non-tEBC; rat-averaged Fisher z: 0.325 vs 0.628),. This disruption was significant across animals (paired t-test in Fisher z space: t(4)=4.392, p**=0.01, n=5 rats) and across session pairs (t(14)=4.805, p<0.001 n=15 days). See Fig. S15 for the corresponding population vector analysis.

This reduction in spatial information arises because neural activity during tEBC is only weakly constrained by spatial location. At the population level, the similarity between population vectors (PVs) corresponding to different tEBC trials was only modestly influenced by spatial position (Fig. 4b-c); though modestly higher for trials occurring within the same quadrant than across different quadrants, this effect was small and inconsistent across animals. A single-cell split-half analysis yielded a similar result, showing only a weak spatial specificity of the temporal activity profiles during tEBC (Fig. 4d). These analyses indicate that neural activity during tEBC is not strongly affected by spatial location.

In contrast, spatial coding was disrupted during the tEBC task (Fig. 4e). The spatial similarity across PVs for matched locations across quadrants was markedly reduced during tEBC periods compared to non-tEBC periods, even when both were compared to the same spatial template derived from non-tEBC activity (Fig. 4f). This reduction was large, consistent across sessions and animals, and highly significant relative to controls based on permutations within a given session (data not shown). A similar effect was found at the single-cell level: for each cell, similarity between its spatial activity and its baseline spatial map was strongly reduced during tEBC (Fig. 4g). In addition, spatial stability within the tEBC condition itself was reduced for each animal, indicating that spatial activity patterns were not only altered relative to baseline but also internally less consistent (Fig. 4h). These results indicate a loss of stable, location-specific activity for individual neurons; a corresponding reduction in the stability of spatially specific activity was observed in population vector analyses (Fig. S15).

These effects were also evident in complementary analyses of the spatial information associated with single-cell activity. At the single-cell level, calcium event rates did not differ between trials whose occupancy centroids fell inside versus outside the cell’s place field (Fig. S18). In addition, neither distance to the place field center during the tEBC period, nor the spatial separation between the centroids of tEBC trials meaningfully predicted calcium event rate variability across tEBC trials (Fig. S19). Consistent with this, spatial mask analyses showed that calcium events during the tEBC period frequently occurred outside place field regions defined during non-tEBC periods (Fig. S20).

Together, these findings reveal a pronounced asymmetry between the representation of the conditioning task and the representation of space in CA1. Neural activity during tEBC is only weakly constrained by spatial location, whereas the tEBC task substantially degrades the spatial structure of activity, both at the single-cell and the population levels. During tEBC, neurons increase their activity and are frequently active outside established place fields, contributing to a degradation of the spatial representation. These results indicate that tEBC recruits a mode of hippocampal processing that is largely distinct from classical spatial coding and can transiently override it.

## Discussion

Here we tested two related and prevailing hypotheses about hippocampal function. First, that the hippocampus primarily represents spatial location. Second, that hippocampal contributions to nonspatial variables is either during immobility or via a change in gain of the place code, e.g. via altered excitability. Previous studies, which mostly used head-fixed subjects, have led to the hypothesis that the hippocampus encodes nonspatial information during immobility^5,6,16,19,31^. Only one study has measured hippocampal selectivity in ambulating rats^38^; however, because task events were not dissociated from spatial location, it remains unclear whether nonspatial signals reflect changes in place cell gain or the emergence of a distinct, nonspatial population code.

Here, we simultaneously measured the activity of many hippocampal CA1 neurons as rats freely moved in a 2D environment while performing a trace conditioning task. Because conditioning task events occurred randomly throughout the environment, this design enabled a direct test of whether task-related activity reflects modulation of spatial coding or a distinct population-level representation.

We found four main results. First, dorsal CA1 calcium event rates were higher during the tEBC period than otherwise, and this increase was not explained by running speed or spatial location. Second, substantially more cells were tuned to the tEBC task than to spatial position. Third, tEBC-related activity was largely independent of spatial location. Fourth, the converse was not true: tEBC engagement disrupted spatial coding, reducing the reliability of place-related activity. These results, obtained at the single cell level, were further confirmed at the population level using population vector analyses.

The tEBC task reorganized hippocampal CA1 activity into a temporally structured population response that was only weakly constrained by spatial location. Although individual neurons exhibited diverse responses to tone, trace, shock, and post-shock epochs, population activity evolved along a consistent trajectory through neural space that was distinct from trajectories during non-tEBC periods of the same duration. This structure was evident at fine temporal resolution, with reliable but distinct population vectors across successive segments of the task. These dynamics are not consistent with a simple gain modulation of ongoing place activity; they reflect the emergence of a task-specific population code.

During the conditioning task, spatial coding was transiently disrupted: place-related calcium events became less reliable, and spatial similarity decreased at both the single-cell and population levels. Together, these findings show that during tEBC, hippocampal coding can be reorganized by task demands into a nonspatial, temporally structured population response that disrupts classical place coding.

Reports that hippocampal activity during nonspatial tasks appears gated by place fields have been interpreted as evidence that nonspatial signals in CA1 are secondary to spatial coding^5,6,38^. However, these place-related responses may reflect the specific behavioral conditions under which those studies were performed. When animals are immobile, or when task events occur at fixed sites or within a limited set of locations, trials are sampled from a narrow spatial range; this causes task-related activity to appear gated by place because spatial variation is not sampled. In contrast, when animals remain mobile and task events are sampled across the entire environment, the tEBC task recruits a temporally organized population response that reduces spatial reliability rather than reinforcing it. We did not observe stable place specific calcium events during tEBC periods, even on tEBC trials preceded by brief immobility. Thus, whether activity during nonspatial tasks appears spatially gated or spatially disruptive reflects behavioral state and spatial sampling, rather than a fixed or obligatory relationship between task-related activity and spatial coding (see Supplementary Box 1 for a detailed comparison). These findings are consistent with a growing body of literature that indicate that a proportion of hippocampal neurons encode nonspatial responses, including social^45,46^, visual^47,48^, and task relevant demands^49^.

Despite their distinct organization, task-related and spatial population structure showed a similar decay with distance. Population vector similarity decreased with increasing temporal separation during tEBC and with increasing spatial separation during exploration; the shapes of these decay functions closely overlapped (Fig 3d). This similarity suggests that task and spatial coding share a common population-level mechanism, even though they are generated by different sensory-motor stimuli and correspond to different dimensions that dominate under different behavioral demands. We thus hypothesize that the trace task recruits an existing population structure rather than introducing a separate representational mode.

Together, these results reveal three core features of hippocampal coding during the tEBC task. First, task-related population activity exceeds spatial coding in magnitude during active behavior. Second, the tEBC task reduces spatial reliability, demonstrating that task coding is not simply layered onto stable place representations. Third, the tEBC task and spatial coding share a common decay with distance, suggesting a shared population mechanism. Apparent disagreements with these observations in the existing literature are likely to arise from differences in which dimension (space or task time) dominates during a behavioral task. Our findings extend the classical view of the hippocampus as a primarily spatial map toward a flexible population code that prioritizes behaviorally relevant representations across time and space.

## Methods

### LEAD CONTACT

Questions and requests for information should be directed to and will be fulfilled by the Lead Contact, Hannah Wirtshafter. This study did not generate new unique reagents.

### DATA AVAILABILITY

The data that support the findings of this study are available from the corresponding author, Hannah Wirtshafter. All code produced for data analysis is available on https://github.com/hsw28/ca_imaging

### EXPERIMENTAL MODEL AND SUBJECT DETAILS

All procedures were performed within Northwestern Institutional Animal Care and Use Committee and NIH guidelines. Five male Long Evans rats (275–325 g) were sourced from Charles River Laboratories, injected with AAV9-GCaMP8m, implanted with a 2-mm GRIN lens, and trained and tested on eyeblink conditioning in a rectangular apparatus (Fig. 1). Animals were individually housed in an animal facility with a 12/12 h light/dark cycle.

## METHOD DETAILS

### GCaMP8 injection, lens implantation, EMG implantation

GCaMP8 injection and lens implantation were completed as reported in Wirtshafter and Disterhoft, 2022, in Wirtshafter and Disterhoft, 2023, and in Wirtshafter, Solla, and Disterhoft, 2024^18,50,51^. Briefly, rats were anesthetized with isoflurane (induction 4%, maintenance 1-2%) and a craniotomy was performed at stereotaxic coordinates Bregma AP −4.00mm, ML 3.00mm. 0.6μL of GCaMP8m (obtained from AddGene, packaged AAV9 of pGP-AAV-syn-jGCaMP8m-WPRE, lot v175525, titer 1.3E+13 GC/mL) was injected over 12 minutes (approximate coordinates Bregma AP −4.00mm, ML 3mm, DV 2.95mm relative to skull); then the syringe was raised 0.2mm and an additional 0.6μl of GCaMP7 was injected. We repeated this process once more and at slightly different coordinates in the craniotomy hole, resulting in 4 total injections.

We then aspirated tissue from the craniotomy site using a vacuum pump and a 25 gauge needle. Tissue was aspirated up to and including the horizontal striations of the corpus collosum. A 2mm GRIN lens (obtained from Go!Foton, CLH lens, 2.00mm diameter, 0.448 pitch, working distance 0.30mm, 550nm wavelength) was then inserted into the craniotomy hole and cemented in place using dental acrylic. Animals were given buprenorphine (0.05mg/kg) and 20mL saline, taken off anesthesia, and allowed to recover in a clean cage placed upon a heat pad.

Six to eight weeks after surgery, animals were again anesthetized with isoflurane and checked for GCaMP expression. If expression was seen, baseplates were attached using UV-curing epoxy and dental acrylic. Electrode implantation to record obicularis oculi electromyographic (EMG) activity occurred in the same surgery as baseplate attachment, as described previously^52,53^. Briefly, a connector containing 5 wires was cemented on the front of the animal’s head: 4 wires were implanted directly above the eye in the surrounding muscle (2 for recording, 2 for electrical stimulation). An additional wire was attached to a connector attached to a ground screw located above the cerebellum; this screw was implanted during lens implantation surgery.

### Behavioral environment and training

A single behavioral apparatus was used in these experiments. The environment was a 78.7cm x 50.8cm unscented rectangular enclosure with wire floor and walls and white lighting. A tether containing a plug to relay the EMG activity and to deliver a shock to the rat’s eye was attached to a the eyeblink connector on the rat’s head. The miniscope was plugged into the cemented baseplate. The miniscope and EMG cords were all attached to a commutator for ease of animal movement.

The CS was a 250ms, 85dB free-field tone (5ms rise-fall time). The US was a 100ms shock directed to the left eye. Shock amount varied per session per animal and was calibrated, if needed, at the end of a training session for the next session’s training. Shock level was deemed appropriate when a shock was met with a firm shake of the animal’s head.

The trace interval was 500ms and the intertrial interval (ITI) was randomized between 30s and 60s, with a 45s average. EMG signals were amplified (5000×) and filtered (100Hz to 5kHz), then digitized at 3kHz and stored by computer.

A conditioned response (CR) was identified as an increase in integrated EMG activity that exceeded the baseline mean amplitude by more than four standard deviations, sustained for a minimum duration of 15ms. Baseline mean amplitude was calculated during the 500ms preceding CS onset. Additionally, the response had to commence at least 50ms after the conditioned stimulus (CS) onset and before the unconditioned stimulus (US) onset.

The animal’s first exposure to each environment was a 38min exploration session, in which the animal was able to freely move and explore the environment without any conditioning (Figure 1a). Animals were then trained in one environment per session, with no more than one session per day, and were considered to have learned the task after reaching criterion (70% CRs in 50 trials) on three consecutive training sessions (termed ‘criterion sessions’).

### Calcium imaging

Calcium imaging was completed as reported in Wirtshafter and Disterhoft, 2022, in Wirtshafter and Disterhoft, 2023, and in Wirtshafer, Solla, and Disterhoft 2024^18,50,51^. Briefly, calcium imaging was done using UCLA V4 Miniscopes^54,55^, assembled with two 3mm diameter, 6mm FL achromat lens used in the objective module and one 4mm diameter, 10mm FL achromat lens used in the emission module.

## QUANTIFICATION AND STATISTICAL ANALYSIS

Unless otherwise noted, means are reported as mean ± standard deviation. All analyses exclude cells with fewer than five calcium events across all trial periods. For comparisons including all animals, averages were computed by first calculating the relevant metric per animal and then averaging across animals, rather than pooling across sessions or cells. The exception is the population-level analyses in Fig. 3, which were computed across all cells. All statistical tests are two sided. All analysis and figure-generation code are available at https://github.com/hsw28/ca_imaging.

### Position and speed analysis

Position was sampled by an overhead camera at 30Hz. Position tracking was done post-recording using DeepLabCut^56^. Position was then converted from pixels to cm and smoothed using a Gaussian filter with σ = 2cm. Speed was calculated by taking the Euclidean distance between the two positions just before and just after the time of interest.

### Video pre-processing and cell identification

Video preprocessing and cell identification were performed as reported in Wirtshafter and Disterhoft, 2022, in Wirtshafter and Disterhoft, 2023, and in Wirtshafter, Solla, and Disterhoft, 2024^18,50,51^. In brief, videos were recorded with Miniscope software at 15frames/second. Video processing was done using CIATAH software^57^. Videos were down sampled in space and normalized by subtracting the mean value of each frame from the frame. Each frame was then normalized using a bandpass FFT filter (70-100cycles/pixel) and motion corrected to a using TurboReg^58^. Videos were then converted to relative florescence (dF/F_0_); F_0_ was the mean over the entire video.

Cells were automatically identified using CIATAH^57^ using CNMF-E^59^. Images were filtered with a gaussian kernel of width 2 pixels and neuron diameter was set at a pixel size of 8. The threshold for merging neurons was set at a calcium trace correlation of 0.65; neurons were merged if their distances were smaller than 4 pixels and they had both highly correlated spatial shapes (correlation>0.8) and small temporal correlations (correlation <0.4).

All cells identified using CNMF-E were then scored as neurons or not neurons by a human scorer. Scoring was also done within CIATAH software in a Matlab GUI. Scoring was done while visualizing, based on the calcium activity trace, average waveform, a montage of the candidate cell’s Ca2+ events, and a maximum projection of all cells on which the candidate cell was highlighted. The local maxima of the relative fluorescence (ΔF/F_0_) of each identified cell were identified as calcium event times.

We recorded a total of 6664 cells across 15 sessions, three sessions per rat (average of 205 ± 51, 339 ± 169, 242 ± 21, 777 ± 121, and 658 ± 213 cells per rat). All analyses included only cells with either a calcium rate of at least 0.05Hz during non-trial periods or an event rate of at least 0.05Hz during trial periods. This resulted in a final dataset of 6192 cells, with an average of 179±34, 328±163, 220±2, 732±130, and 603±203 cells per animal.

### Task modulation analysis (conditioning vs intertrial)

Trial period activity was defined as the 0s–2s window following CS onset. All other time points were considered non-trial periods unless otherwise noted. Task modulation was quantified as the ratio of mean activity during the trial period to mean activity during the intertrial interval. Significance was assessed by comparing the observed modulation value for each neuron to a null distribution generated by permuting trial and intertrial labels 500 times.

### Calcium event rate comparisons

We computed per neuron calcium event rates during the trial window and during randomly sampled non-trial windows of the same duration. We compared the distribution of calcium event rates across conditions using a two-sample Kolmogorov–Smirnov (KS) test and a paired t-test. Population-level differences in mean event rate were visualized using histograms and bar plots of mean ± SEM.

### Modulation clustering by epoch

To identify neurons with selective modulation during different phases of the trace conditioning period, we computed calcium event rate changes relative to baseline in four task epochs: conditioned stimulus (CS; 0–250ms after tone onset), trace interval (250ms–750ms), unconditioned stimulus (US; 750ms–850ms), and a post-US period (850ms–2000ms). Baseline activity was computed from activity during a pretrial window (−2s to 0s relative to CS onset) for each neuron and for each trial. For each neuron, we computed a modulation index for each epoch as:

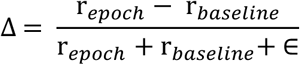

where:

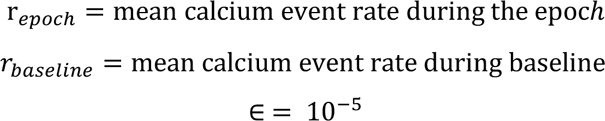

Because CS (250ms) and US (100ms) are short relative to our 133ms frame, very low counts inflate variance and can saturate the index Δ. To ensure well estimated rates, we only included cells with ≥5 total events across all trials in the 2s task window OR with ≥5 baseline events during the 2s preceding the task window. This criterion is direction agnostic (does not favor excitation) and was chosen based on stability diagnostics. As a robustness check, we repeated the analysis using PSTHs with data centered around the mean and obtained the same directional effects (Fig. S6).

### Speed modulation within and across trials

To test whether calcium event rates were modulated by running speed at the trial level, we calculated the Pearson correlation between mean speed and mean calcium event rate over the 2s tEBC window following CS onset across trials for each neuron. Mean speed was computed within the 2s tEBC window. Neurons with fewer than five total calcium events across all trials were excluded. To assess significance, we applied a false discovery rate (FDR) correction (q = 0.05) to the set of correlation p-values for each animal. Additionally, we compared the observed correlations to a null distribution obtained by randomly permuting trial speeds 500 times and recalculating the correlation for each permutation. Neurons whose observed correlation exceeded the 95th percentile of the null distribution were considered significantly modulated by running speed.

To assess whether the rate of calcium events was modulated by time dependent speed fluctuations within trials, we binned the tEBC period into 15 bins of duration 0.133s and computed the Pearson correlation between speed and calcium event rate within each bin for each trial. For each neuron, these binned correlations were averaged across trials. Neurons with fewer than five total calcium events across all trials were excluded. Statistical significance was assessed using an FDR correction (q = 0.05) across the population. We generated a null distribution by randomly permuting the speed values within bins within each trial 500 times and recalculating the correlations for each permutation. The fraction of neurons whose correlations exceeded the 95th percentile of the null distribution was used as an estimate of the false positive rate.

### Speed-matched and acceleration-matched comparison of tEBC and non-tEBC activity

We asked whether hippocampal neurons changed their calcium event rate during the tEBC period relative to non-tEBC periods of the same duration while controlling for running speed. The tEBC period was defined as the 2s window beginning at CS onset. Non-tEBC periods were defined as all frames outside the excluded task windows. Running speed was computed framewise from consecutive positions as the Euclidean displacement divided by elapsed time.

To control for speed, tEBC and non-tEBC frames were grouped into 2 cm/s speed bins using common bin edges defined from the range of speeds observed during the tEBC period. For each speed bin, we pooled all tEBC frames in that speed bin across the session; separately, we pooled all eligible non-tEBC frames whose speed matched the same speed bin. For each cell, calcium event rate in that speed bin was computed as the total number of events divided by the total pooled duration; differences in the amount of available non-tEBC data affected precision but not the event rate estimate. This procedure allowed us to compare trial and non-trial activity within matched speed bins, rather than by pairing individual trial windows to individual non-trial windows. Bins were included only if both conditions contributed nonzero occupancy, and cells were included if they had at least one such matched bin. For each cell, differences between tEBC and non-tEBC calcium event rates were computed within each matched bin, and the mean of these per-bin differences (equal weighting across bins) was used as a summary effect size.

At the population level, significance was assessed with a sign-flip permutation test on the mean trial to non-trial differences per cell. On each permutation, the sign of the difference observed for each cell was randomly flipped, and the population mean was recomputed. The resulting null distribution represents the expectation of no consistent directional shift across cells between tEBC and non-tEBC periods. The observed mean difference across the population of cells was then compared to this null distribution to obtain a two-sided permutation p-value.

The same procedure was used for acceleration matching, except that acceleration was computed as the first derivative of speed and grouped into bins of 3 cm/s².

### Population vector similarity

We quantified the structure of population activity during tEBC using a repeated random sampling analysis of population vectors based on the calcium event rate of each neuron. For each session, matrices of calcium event rates were constructed for all tEBC trials in that session by computing the calcium event rate of each neuron during each 2s tEBC period. This matrix contains as many rows as neurons and as many columns as tEBC trials during the session. To assess similarity across tEBC trials, we randomly partitioned the available tEBC trials into two non-overlapping groups of 25 trials each and averaged calcium event rates for each neuron across trials within each group to obtain two population vectors (PVs); the components of these vectors are calcium event rates for each neuron over the same set of neurons within the session. Similarity between these two trial averaged PVs was quantified as the Pearson correlation across neurons.

To compare tEBC to non-tEBC activity, we sampled 2s windows that did not overlap with any tEBC windows. For analyses controlling for locomotor confounds, speed matched non-tEBC windows were selected by matching the speed profile within each tEBC trial window to that of candidate non-tEBC windows by comparing the within-window speed profile of each tEBC trial to candidate non-tEBC windows using the speed-binning procedure described above.

This procedure yielded five within-session similarity measures: tEBC–tEBC, non-tEBC–non-tEBC, tEBC–non-tEBC, tEBC–speed matched non-tEBC, and speed matched non-tEBC–speed matched non-tEBC. For each metric, a random resampling was repeated 500 times per session to generate similarity distributions; session means were used for statistical comparisons. All similarity analyses were performed within session, and sessions were weighted equally in pooled summaries (Fig. 3a).

For an analysis of the temporal structure within a tEBC period (Fig. 3b, S12), the 2s trial window was subdivided into 15 equal temporal bins defined on the imaging-frame time base. In each bin, a PV followed from computing the mean calcium event rate for each neuron across the imaging frames included in the bin. Pairwise Pearson similarity across bins produced a temporal similarity matrix per session. Lag–similarity curves were generated by averaging entries along successive matrix diagonals. The temporal structure associated with the segmentation of a tEBC period across epochs followed from grouping bins corresponding to CS, trace, US, and post-US epochs (Fig. 3b, S11–S12).

For the analysis of the spatial structure of the neural activity during free running non-tEBC periods (Fig. 3c, S13–S14), frames corresponding to non-tEBC while running at speeds ≥ 4 cm/s were grouped into 15 equal occupancy spatial bins. A PV for each spatial bin was computed as the mean calcium event rate across imaging frames in that bin, using the same per-cell mean-centering normalization applied to temporal PVs. Pairwise Pearson similarities across spatial bins resulted in a spatial similarity matrix from which lag–similarity curves were derived.

The reliability of temporal and spatial structure was assessed using even–odd split-half analyses, in which trials or frames were divided into even- and odd-indexed subsets (Fig. S12, S14). Lag curves from each split were correlated to confirm consistency within each session.

### Lag–similarity analysis

For the lag similarity analysis (Fig. 3d), temporal and spatial similarity matrices were reduced to lag similarity curves by averaging entries along their diagonals. We generated null curves by permuting the joint row/column order of each similarity matrix 500 times and recomputing lag curves.

To compare temporal and spatial organization directly, lag-similarity curves were centered around the mean and their correspondence quantified using Pearson correlation and RMSE across bins. Differences in overall magnitude between temporal and spatial curves were tested using permutation tests on the difference in area under the curve (ΔAUC). Analyses were repeated across multiple bin numbers (4–14 bins) to verify robustness.

### Spatial interference in task coding: population-vector (PV) and single cell analysis

In each session, calcium event activity was aligned to imaging frames and analyzed within a fixed 2D spatial grid (4 quadrants in a 2 × 2 grid), and neural activity was measured as calcium event rates per cell. Prior to downstream analyses, the activity of each cell was centered around the mean activity of that cell over the duration of a session, so that similarity metrics reflected relative modulation patterns rather than global differences in calcium event rate. The 2s tEBC window was subdivided into 15 equal temporal bins. For each trial, temporal bin, and spatial bin, a population vector (PV) was computed as the activity of the recorded cells, averaged across the frames in the corresponding temporal–spatial intersection of bins.

To quantify spatial structure during tEBC, PV similarity was assessed using a split-half procedure. Within each temporal bin and spatial bin, the trials in a session were randomly divided into two non-overlapping halves; PV templates were computed by averaging separately within trials in each half. SAME-space similarity was defined as the Pearson correlation between templates from the same spatial bin across trial halves, while DIFFERENT-space similarity was defined as the Pearson correlation between templates from different spatial bins across trial halves. Correlations were Fisher-z transformed and averaged across spatial bins and temporal bins to yield similarity measures at the session level.

Split-half resampling was repeated 50 times per session, and results were averaged across resamples to obtain stable estimates at the session level. These values were then averaged across sessions for each animal. Statistical significance was assessed using paired comparisons in Fisher-z space at both the session level (n = 15 sessions, 3 sessions per rat) and the animal level (n = 5 rats). Values shown in Figs. 4b-c are inverse Fisher-transformed^60^ (tanh) for interpretability.

For single cell analyses, the same temporal and spatial binning and split-half procedure were used. For each neuron, temporal activity vectors across bins were computed separately for each spatial bin and split half. SAME-place correlations were defined as correlations between temporal vectors from the same spatial bin across halves, with SAME-bin and DIFFERENT-bin controls defined analogously. Fisher-z–transformed correlations were averaged across neurons to yield session-level and animal-level similarity measures.

### Task interference in spatial coding: population-vector (PV) and single cell analysis

In each session, calcium event activity was aligned to imaging frames and analyzed within a fixed 2D spatial grid (4 quadrants in a 2 × 2 grid), and neural activity was measured as calcium event rates per cell. Control frames corresponding to non-tEBC periods excluded all frames within a 2s window following CS onset plus an optional buffer extending beyond the tEBC window; analyses optionally applied a running-speed threshold. Control frames were randomly split into two halves. For each spatial bin, a population vector (PV) was computed as the activity of the recorded cells, averaged across frames in that bin. Prior to constructing rate maps and population vectors, the activity of each cell was centered around the mean activity of that cell over the duration of a session, to remove cell specific baseline activity rate offsets so that similarity metrics reflected relative spatial modulation patterns rather than global calcium event rate differences.

One half of the control data was used to construct a spatial template PV for each bin. Non-tEBC spatial reliability was quantified as the Pearson correlation between the template PV and the PV computed from the half held out for control (split-half reliability).

To quantify spatial coding during the tEBC, PVs were computed from frames within the tEBC window and grouped by spatial bin. For each spatial bin, the task PV was correlated with the corresponding control template PV from the same session. Correlations were Fisher-z transformed and averaged across bins to yield a spatial similarity measure at the session level; these were then averaged across sessions for each animal (Fig. 4e).

For single cell analyses (Fig. 4f), the same framework was applied at the level of individual neurons: for each cell, spatial calcium event rate maps were constructed across bins, and reliability was quantified as the correlation of each cell’s spatial map across conditions or across half-split trials, followed by Fisher-z averaging across cells.

To reduce sampling variability, the control half-split and the template construction were repeated 50 times with random resampling, and Fisher-z values were averaged across resamples prior to statistical testing. Task effects were assessed by comparing tEBC versus non-tEBC Fisher-z values using a within-session sign-flip permutation test applied to differences across bins, yielding two-sided p-values that preserve spatial structure while randomizing task labels. Values shown in figures reflect the inverse Fisher transform (tanh) of mean z values for interpretability.

For spatial stability analysis (Fig. 4g), for each animal and recording day, we quantified the stability of coarse spatial calcium event rate patterns on a fixed 2×2 spatial grid that divides the arena in four quadrants (2 x 2 grid). Different baselines in calcium event rates for different cells introduce variable offsets unrelated to spatial patterning; rate maps were thus optionally normalized in each session by centering each cell around its mean using pooled finite map entries across all bins and windows, applied identically to tEBC and non-tEBC maps of calcium event rates.

We estimated spatial stability using split-half correlations. For each session and condition (tEBC vs non-tEBC), windows were randomly split into two halves and averaged within each half to obtain independent estimates of the spatial calcium event rate pattern; this procedure was repeated 50 times with different random splits. We report two complementary stability metrics, each computed as Pearson correlation followed by Fisher z-transform and averaging in z-space. The stability of single cell maps was computed per cell as the correlation between mean calcium event rate vectors from the four bins, split in half, using pairwise valid bins (minimum two valid bins), and then averaged across cells and split-half repetitions. The stability of population vectors (PV) was computed within each spatial bin as the correlation across cells between the two halves population vectors for that bin, using pairwise valid cells (minimum 10 valid cells), and then averaged across bins and split-half repetitions. Pearson correlation coefficients were Fisher z-transformed prior to averaging and statistical testing. Values shown in figures reflect the inverse Fisher transform (tanh) of mean z values for interpretability. tEBC vs non-tEBC differences were assessed with paired t-tests performed on Fisher z values either at the animal level (mean across sessions within each animal) or pooled across session pairs.

### Place cell identification and mutual information

Place cells were identified using mutual information (MI) computed when the animals were running at speeds ≥ 4cm/s. The MI was computed for all cells; there was no calcium event rate criterion for cells to be included. To be considered significant, the computed (MI) must be greater than 95% of MI scores computed 500 times from shuffled positions^61^.

To compute the MI for each cell, the training environment was divided into 2.5cm x 2.5cm spatial bins. The arena is thus divided in *N* bins, labeled by an index *i*, 1 ≤ *i* ≤ *N*. At each time frame, we determine the bin *i* where the rat is located, and record whether a calcium event occurred (*k* = 1) or not (*k* = 0) at that bin at that time. The mutual information between location and cell activity is given by^62–64^:

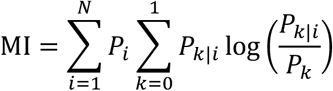

where

*P_i_* = the probability of occupancy of bin *i*

*P_k|i_* = conditional probability of observing a calcium event (*k* = 1) or not (*k* = 0) at bin *i*

*P_k_* = the event probability, given by

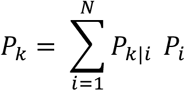

To evaluate the conditional probability *P_k|i_* we consider only those frames when the rat occupies bin *i*, and count *n*(*k* = 0, *i*), the number of frames when the rat was in bin *i* and there was no calcium event, and *n*(*k* = 1, *i*), the number of frames when the rat was in bin *i* and there was a calcium event. The es|mated condi|onal probabili|es follow:

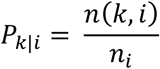

with 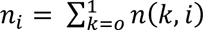, the occupancy count of bin *i*. The estimated occupancy probability *P_i_* is the ratio between *n_i_* and the total number of frames.

To identify place cells, Mutual Information was computed during periods of free movement; only non-tEBC data that excludes the 2s window following each CS onset was used. To quantify how calcium events during tEBC influence spatial information, we recomputed MI including the 2s tEBC windows for all trials within a session. Including the conditioning task periods consistently reduced MI (Fig 4a) in every animal, indicating that neural activity during tEBC trials tended to be nonspatial.

To ensure that this reduction was not driven solely by loss of event counts, we performed an event matched control: for each neuron we removed the same number of calcium events as those that took place during the 2s tEBC windows; the removed calcium events were chosen at random among events that occurred during non-tEBC periods with the animal moving at speeds ≥ 4cm/s. The control MI was recomputed using this randomly reduced set of calcium events.

### Calcium event rate variability by location in place field and distance

The spatial position of the animal during each tEBC trial was assigned using the centroid of the occupancy mode within the 2s tEBC window. Specifically, we constructed a 2D histogram of the animal’s position during each trial using 2.5cm × 2.5cm spatial bins and defined the trial centroid as the center of the bin with the highest occupancy. The histogram grid was aligned to the spatial grid used for rate maps so that trial centroids and place-field masks were defined in a common coordinate frame.

For each cell, we computed the calcium event rate during the 2s tEBC window on each trial and assigned the corresponding trial centroid. To assess whether activity during tEBC depended on spatial position relative to the cell’s place field, we compared the mean calcium event rates of trials whose centroids fell inside versus outside the place field defined from activity during non-tEBC periods (Fig. S18). Comparisons were performed at the single-cell level using paired, two-sided t-tests.

We next examined whether tEBC activity varied as a function of distance from the centroid assigned to the tEBC trial to the place-field center (Fig. S19). For each trial, we computed the Euclidean distance between the trial centroid and the center of the place field, defined as the centroid of spatial bins exceeding a threshold of 1σ above the mean in the non-tEBC rate map. Trial calcium event rates were then regressed against this distance, and Pearson correlation coefficients were computed for each cell and at the population level.

Finally, to test whether spatial proximity between trials predicted similarity in activity during tEBC independently of place-field membership, we computed all pairwise comparisons of trials for each cell (Fig. S20). For each pair of tEBC trials we calculated the Euclidean distance between their centroids and the normalized difference in calcium event rates, defined as |Δ rate| divided by the mean of the two rates. Pearson correlations between spatial separation and calcium event rate variability were computed for each cell and across the pooled dataset.

### Rate map masking and tEBC vs. non-tEB calcium event rate comparison

To assess whether calcium events during tEBC periods occurred outside of typical place fields, we created spatial masks based on rate maps computed separately for tEBC and non-tEBC periods. We computed 2D calcium event rate maps using spatial histograms normalized by occupancy. The bin size was 2.5cm x 2.5cm; both event count and occupancy were smoothed with a 2D gaussian kernel with σ = 0.5 bin = 1.25cm. For each condition, tEBC or non-tEBC, we defined a binary spatial mask by thresholding the calcium event rate map at one standard deviations above the mean rate across spatial bins. These high-rate masks represent putative place fields during each condition. Next, we computed the fraction of calcium events that occurred within each mask during both tEBC and non-tEBC periods. This yielded four calcium event rate values per neuron: calcium event rate during tEBC in a tEBC-defined mask, calcium event rate during non-tEBC in a tEBC-defined mask, calcium event during tEBC in a non-tEBC-defined mask, and calcium event rate during non-tEBC in a non-tEBC-defined mask. These values were compared using paired two-sided t-tests to assess whether activity in each spatial mask was significantly modulated by behavioral state (Fig. S21).

Comparisons of calcium event rates restricted by the masks allowed us to assess whether tEBC calcium events occurred preferentially in zones of high activity during tEBC or zones of high activity during non-tEBC: the place field zones.

## Acknowledgements

This work was supported by an NIA T32 (T32-AG020506/AG/NIA), an NIA R37 (R37-AG008796/AG/NIA), an NINDS R01 (R01 NS113804/NS/NINDS), and a K99 award (K99 MH135062). We would like to thank all members of the Disterhoft lab, especially Mackenzie Kneisly.

## Author contributions

Investigation, H.S.W.; Formal Analysis, H.S.W.; Writing – Original Draft, H.S.W.; Writing – Review & Editing, H.S.W., M.R.M, S.A.S., J.F.D.; Funding: H.S.W, J.F.D., Supervision, M.R.M, S.A.S., J.F.D.

