## Supplemental Captions for "Hippocampal conditioning code dominates and disrupts the place code"

**Supplementary Box 1**

| Prior task paradigms | This study |
|---|---|
| Fixed stimulus location or trials that tend to happen at specific locations such as endpoints[38] | Random trial locations (Fig 1) |
| Long CS or repeated US[39] | Brief CS-trace-US structure (Fig 1) |
| Reward linked to endpoints[38] or rewarded fields[39] | No reward (Fig 1) |
| Non-hippocampal dependent task (e.g. no trace period)[39] | Hippocampal dependent task[30] |
| Freezing at trials[39] | No freezing at trials (Fig 2) |
| Electrical artifact during US limits reliable recording of hippocampal activity during shock[38] | Calcium imaging permits continuous measurement through CS, trace, US, and post-US epochs |
| **Observed outcome:** Task responses appear gated by place fields when spatial and task signals align | **Observed outcome:** Temporally structured task ensemble (Figs. 2–3) accompanied by reduced spatial stability (Fig. 4) |
| Whether hippocampal task responses appear spatially gated or spatially disruptive depends on behavioral state and spatial sampling, reflecting flexible coding rather than a fixed representation of only space[12]. | |

**Box 1 | Behavioral and methodological factors that determine whether hippocampal task responses appear place bound.**
Prior reports of place gated responses during nonspatial tasks could arise from restricted spatial sampling, immobility, reward structure, or limited access to brief task epochs. When these constraints are removed, the nonspatial task recruits a temporally structured pattern of population activity that reduces spatial stability.

**Supplement captions**

**S1.** Learning performance across the five rats. Bars indicate the average percent of trials with a CR during criterion sessions; each dot represents one session. Dashed red line indicates the 70% learning threshold.

**S2.** Example task-modulated cells. Left to right: trial-aligned rasters, event rate histograms, and average calcium signals (mean ± s.e.m.) for individual cells. Vertical dashed line indicates CS onset; shaded region marks the conditioning window (0–2 s post-CS). Cells exhibited heterogeneous responses across the CS, trace interval, US, and post-US period.

**S3.** Top row left: Mean calcium event rate during the trial period (0.10 ± 0.02 Hz) was significantly higher than during all non-trial periods (0.07 ± 0.01 Hz), as well as non-trial periods restricted to speeds above 4cm/s (0.08± 0.02 Hz) (both paired t-tests, ***$p < 0.001$). Distribution of firing rates across cells also differed significantly between task and non-task periods with speed > 4.0cm/s (two-sample Kolmogorov–Smirnov (KS) test, $p < 0.001$), and for task periods and non-task periods at any speed > 4.0cm/s (two-sample KS test, $p < 0.001$, data not shown). Below: same data but for all five rats.

**S4.** Non-trial vs. trial comparisons of firing and information metrics, plotted per cell for each of five rats (rows 1–5) and pooled across all animals (bottom row). Columns show: (1) normalized firing rate (each cell's rate divided by its session mean), (2) raw firing rate in Hz. In each subplot, the dashed diagonal is unity ($y = x$). All rats exhibit strong, highly significant positive coupling between pre- and task-epoch for both normalized and raw rates.

**S5.** Same as Figure 2c but restricted to neurons identified as significant place cells (spatial MI > 95% of shuffled control). Left: For each task epoch (Tone/CS, Trace, Shock/US, Post), bars show the number of cells significantly modulated relative to non-trial activity. Stacks partition each epoch's modulated population into cells modulated only in that epoch (blue) or additionally in +1 (green), +2 (gray), or +3 (purple) other epochs. This visualizes how strongly modulation is restricted to a specific epoch versus shared across multiple task periods. Right: The same counts normalized by epoch duration (cells per second of the epoch window). This compensates for large differences in window length (e.g., the short 100 ms shock epoch vs. the longer post epoch) and reflects the density of modulation per unit time.The pattern of task-epoch selectivity closely matched that of the full population, with no significant difference in proportions between all cells and place cells (paired t-test, $p > 0.05$; KS test $p > 0.05$).

**S6.** Same as Figure 2e but expanded to show all animals individually. Top row indicates average EMG and speed during the trial time for each animal, shaded areas denote ± 1 SEM.. Middle row displays PSTHs for individual animals. Bottom row shows the across-animal mean after de-meaning each cell's baseline activity. Each trace shows the average firing dynamics of CA1 neurons during trace conditioning, binned at 133 ms from 0.5s before to 4s after tone onset. Tone onset is indicated by a dotted line. The mean centered averages highlight temporally structured responses to the CS, trace, and US epochs that are preserved across animals.

**S7.** Running speed during the 2 seconds post CS onset (trial, in blue) versus the entire matched pre-trial period (pre-trial, in purple), for each animal. Black bars indicate means, black circles indicated medians. For visualization, violins are truncated at the 99th percentile; values above the cap are clipped. Means/medians are computed on the full data.*$p < 0.05$, paired t-test.

**S8.** Top: Across trial (trial level) correlations between mean speed and mean calcium activity. Only a small fraction of neurons appeared significantly speed-modulated, and none exceeded significance relative to a shuffled control. Left: Mean Pearson correlation coefficients (r ± SEM) between speed and event rate were -0.029± 0.165, −0.023 ± 0.174, 0.003 ± 0.151, −0.047 ± 0.222, and 0.013 ± 0.179 for rats 1–5, respectively. Center left: Distribution of r-values across all cells. Center right: Percentage of significantly modulated neurons per rat: 0.0%, 2.2%, 0.0%, 7.7%, and 0.4% ($p < 0.05$, FDR-corrected). Far right: Percentage of significantly modulated cells relative to shuffled controls per animal: 3.7%, 0.0%, 0.0%, 8.7%, and 3.7%. Bottom: Within-trial correlations between speed and calcium activity. Layout as in (b), but for correlations computed within each trial. Left: Mean Pearson r-values (± SEM) were 0.0501± 0.120, 0.075 ± 0.132, 0.061 ± 0.108, 0.057 ± 0.113, and 0.028 ± 0.111 for rats 1–5. Center left: Distribution of r-values across cells. Center right: Percentage of significantly modulated cells was 0.0%, 2.3%, 0.0%, 0.5%, and 1.0%, respectively ($p < 0.05$, FDR-corrected). Far right: No animals showed significant modulation relative to shuffled controls (0% in all cases).

**S9.** Same as Figure 2h but matched for acceleration instead of speed. Calcium event rates for the population were compared between task trials and inter-trial epochs matched for mean acceleration. Each distribution shows the fold change in event rate (trial / non-trial, acceleration-matched) for all cells, averaged per rat. Across all animals, neurons exhibited significantly higher firing during trials than during acceleration-matched controls (all permutation test $p < 0.001$). These results confirm that task-related rate increases are not explained by differences in movement.

**S10.** Same as Figure 2i but matched for acceleration instead of speed. Calcium event rates for single cells were compared between task trials and inter-trial epochs matched for mean acceleration. Each distribution shows the fold change in event rate (trial / non-trial, acceleration-matched) for all cells, averaged per rat. Across all animals, neurons exhibited significantly higher firing during trials than during acceleration-matched controls (all permutation test $p < 0.001$). These results confirm that task-related rate increases are not explained by differences in movement.

**S11.** Same as Figure 3b but displayed for each rat individually. Heatmaps show Pearson similarity between split-averaged population vectors (0–2 s after CS onset, divided into 15 bins). Each rat shows a pronounced diagonal and banded off-diagonals, indicating that ensemble states are most similar at nearby times and diverge progressively with temporal lag. Within-epoch similarity exceeds between-epoch similarity for all animals (paired t-tests $p < 0.001$), confirming consistent temporal organization of CA1 population dynamics.

**S12.** Left: Population-vector (PV) correlation matrices comparing even vs. odd trial blocks in successive 0.133 ms bins after US onset, pooled across rats. Each cell is demeaned (has its own mean across all bins×trials subtracted). The diagonal reflects within-bin similarity, indexing the stability of ensemble activity at each time point. Right: Diagonal correlation values (r) as a function of time since US. Top row: All post-trial bins included, regardless of running speed. Bottom row: Analysis restricted to bins with mean speed > 4 cm/s. Both approaches show that ensemble correlations decay gradually but remain elevated for several seconds after the US. See also figure S22.

**S13.** Same as Figure 3c but for each rat separately. Heatmaps depict pairwise Pearson correlations between population vectors computed from 15 equal-occupancy spatial bins during non-trial running (speed ≥ 4 cm/s). Each rat shows a clear spatial distance-decay pattern, with high similarity along the diagonal and decreasing correlation with inter-bin distance. Adjacent bins are significantly more similar than non-adjacent bins (paired t-tests $p < 0.001$ for all animals), confirming the reliability of the distance-structured spatial code.

**S14.** To verify that the spatial distance-decay was not driven by sampling idiosyncrasies, we computed split-half (even/odd) population vectors within each spatial bin during non-task running and correlated them across bin pairs. The pooled A×B bin matrix showed a strong diagonal and banded off-diagonals, and the mean similarity declined smoothly with inter-bin distance.

**S15.** Population-vector spatial stability is reduced during task epoch. Population vectors (PVs) were constructed from mean activity across spatial bins (2×2 grid) and spatial stability was quantified as the Pearson correlation between PVs computed from split halves of the data (repeated random splits). Points show per-animal means (bold colored lines); lighter lines show individual days; bars show across-animal mean ± SEM. Population-level spatial similarity was higher during non-task than task epochs (mean r: 0.143 vs 0.011; rat-averaged Fisher z: 0.144 vs 0.011). This difference was significant across animals (paired t-test in Fisher z space: $t(4)=3.661$, $p<0.05$) and across pooled day pairs ($t(14)=5.483$, $p<0.001$).

**S16.** Spatial mutual information across cells (MI), computed using behavior outside of trial periods. Each point represents a cell; black lines indicate medians. Across all animals and sessions, the average spatial MI was 1.75±0.23, with averages for individual rats of 1.53 ± 0.63, 1.84±0.55, 2.10±0.79, 1.59±0.63, and 1.67±0.62, and medians of 1.42, 1.74, 1.97, 1.47 and 1.56. When compared to a shuffled distribution, 20.6±3.9% of cells were significantly spatially modulation.

**S17.** Same as 4a, shown separately for each rat. The increase in MI following removal of task-period activity was observed in all animals, while removal of random or speed-matched events produced minimal effects.

**S18.** Paired comparison of per-cell mean firing rates during the CS→2 s task window when trial centroids fell inside versus outside of each cell's non-task-defined place field. Each line connects paired values from a single cell. Left column is individual animals, right column is all animals where statistics reflect a paired two-sided t-test across all animals ($p>0.05$, $t=-0.12$, $n=5075$).).

**S19.** Left: Trial firing rate vs. distance to a cell's place-field (PF) center. For each trial we computed the animal's occupancy-mode centroid during the CS→+2 s window and measured its Euclidean distance to the center of the cell's PF defined from non-task running ($> \mu + 1\sigma$; PF center = centroid of suprathreshold bins). Panels 1–5 show individual rats (rats 1–5); panel 6

shows the pooled data across all rats. Black line, least-squares fit; panel titles report Pearson r, two-sided p, and sample size (trials). Correlations were numerically tiny across animals and pooled ($r \approx 0.01$; $p < 0.05$ due to large N), indicating no biologically meaningful dependence of rate on distance to the PF center. For visualization, a random subsample of points is shown; all data were used for statistics.
Right: Trial-to-trial variability vs. spatial separation of trial centroids. For each cell we formed all unordered trial pairs and computed normalized variability as $|\Delta\text{rate}|/\text{mean}$. The x-axis is the Euclidean distance between the two trials' occupancy-mode centroids (same centroid method as Fig. S16). Panels 1–5 show individual rats (rats 1–5); panel 6 shows the pooled data across all rats. Black line, least-squares fit; panel titles report Pearson r, two-sided p, and sample size (pairs). Correlations were again extremely small (pooled $r \approx 0.01$; $p < 0.05$ with $n \approx 7.6\times10^6$ pairs), reflecting statistical significance driven by sample size rather than effect magnitude. Together with Fig. S16, these results indicate that neither distance to the PF center nor spatial separation between trials meaningfully explains trial-to-trial variability in task-related firing.

**S20.** Spatial masks confirmed that cells frequently had calcium events outside their place fields during the trace task. a. Example spatial masks derived from non-task (left) and task (right) calcium rate maps. Masks highlight regions with activity >1 standard deviations above the mean, used to define high activity zones during each period (these zones represent putative place fields during non-task times). b. Per-cell comparison of out-of-trial vs in-trial calcium event rates. Each scatter shows one neuron's event rate outside versus inside the mask, with the dashed diagonal marking equality ($y = x$) and the solid red line the least-squares fit. Pearson's r and p-value are indicated in each panel. Left: Spatial mask (events inside vs outside each cell's putative place field, determined by mean event rate + 1 standard deviation during non task period). Right: Task-defined mask: events inside vs outside the high event rate 0–2 s CS window. High event rate was determined as mean rate during the task + 1 standard deviation. See also S11 c. Average calcium event rates within the task-defined spatial mask during task and non-task periods. All five rats showed significantly higher activity in task-defined regions during trials (***$p < 0.001$). Right: Average event rates within the non-task-defined spatial mask during task and non-task periods. Two of five rats showed decreased calcium events in non-task (putative place field) zones during trials, suggesting that task period calcium event patterns may supersede place cell event patterns (***$p < 0.001$, **$p<0.01$).
