## Supplemental figures for "Hippocampal conditioning code dominates and disrupts the place code"

### Fig S1

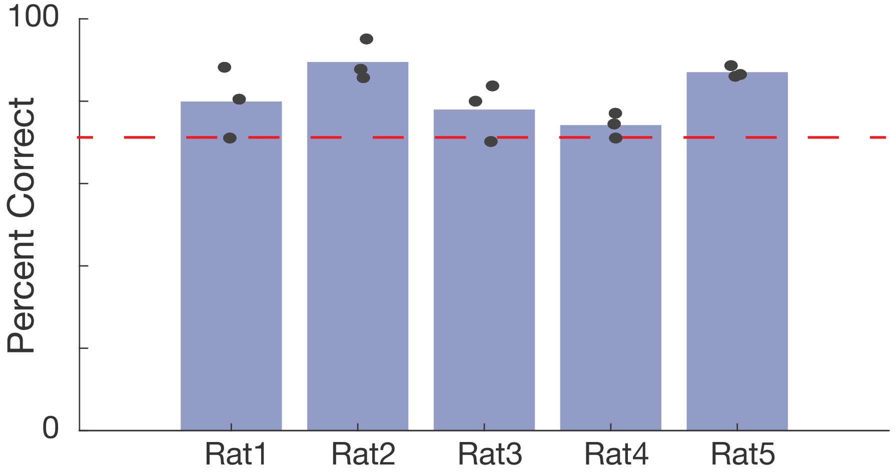

### Fig S2

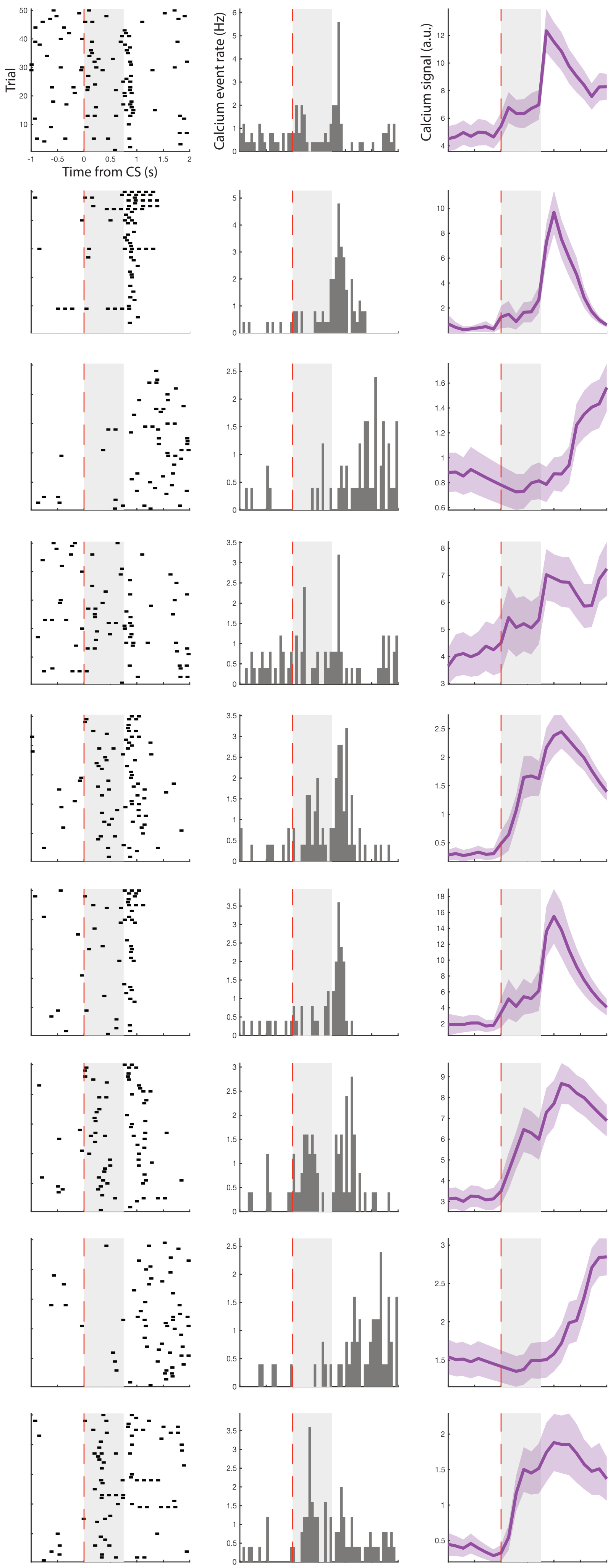

### Fig S3

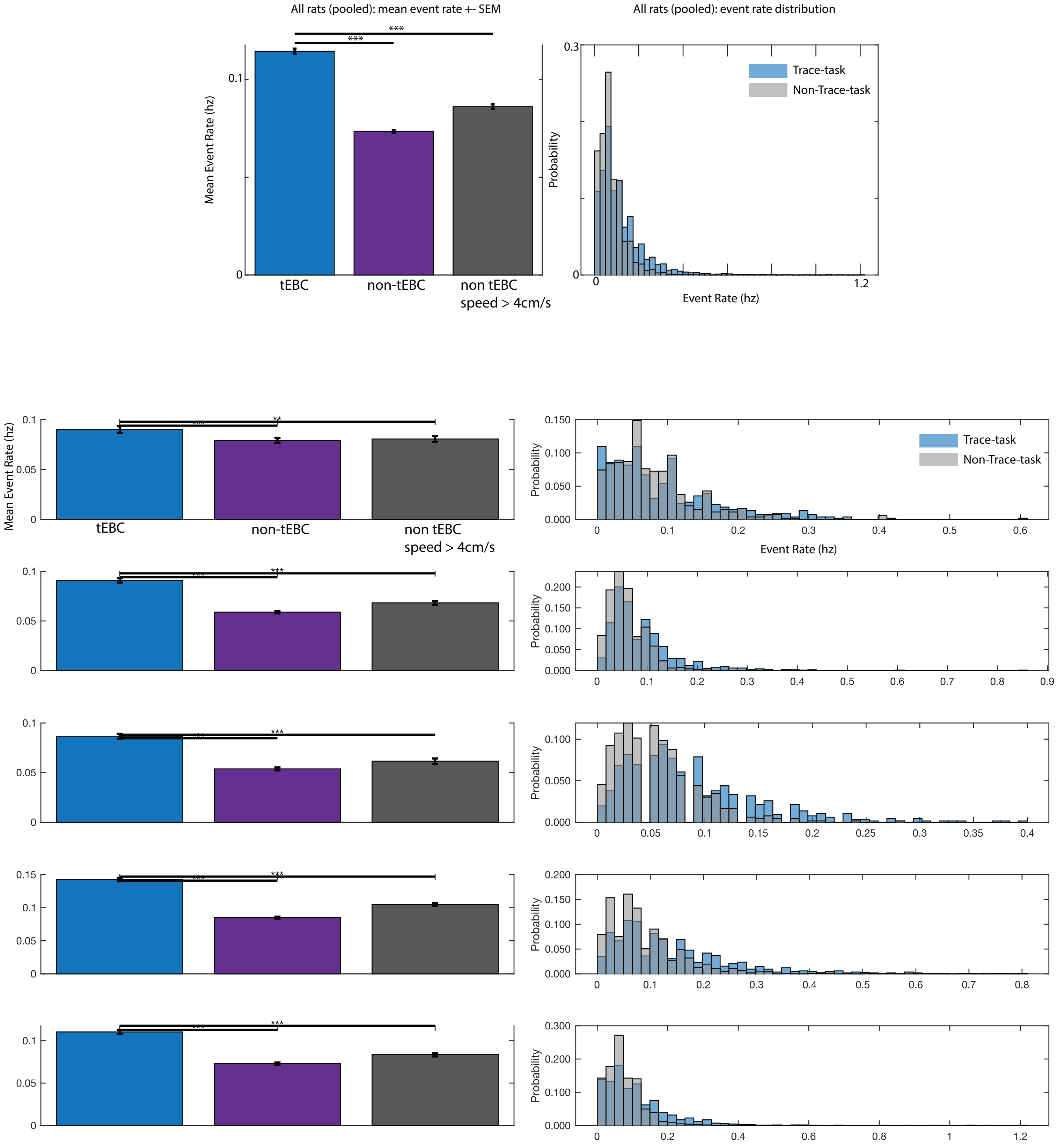

All rats (pooled): mean event rate +- SEM
All rats (pooled): event rate distribution
Mean Event Rate (hz)
tEBC
non-tEBC
non tEBC
speed > 4cm/s
Probability
Event Rate (hz)
Trace-task
Non-Trace-task

### Fig S4

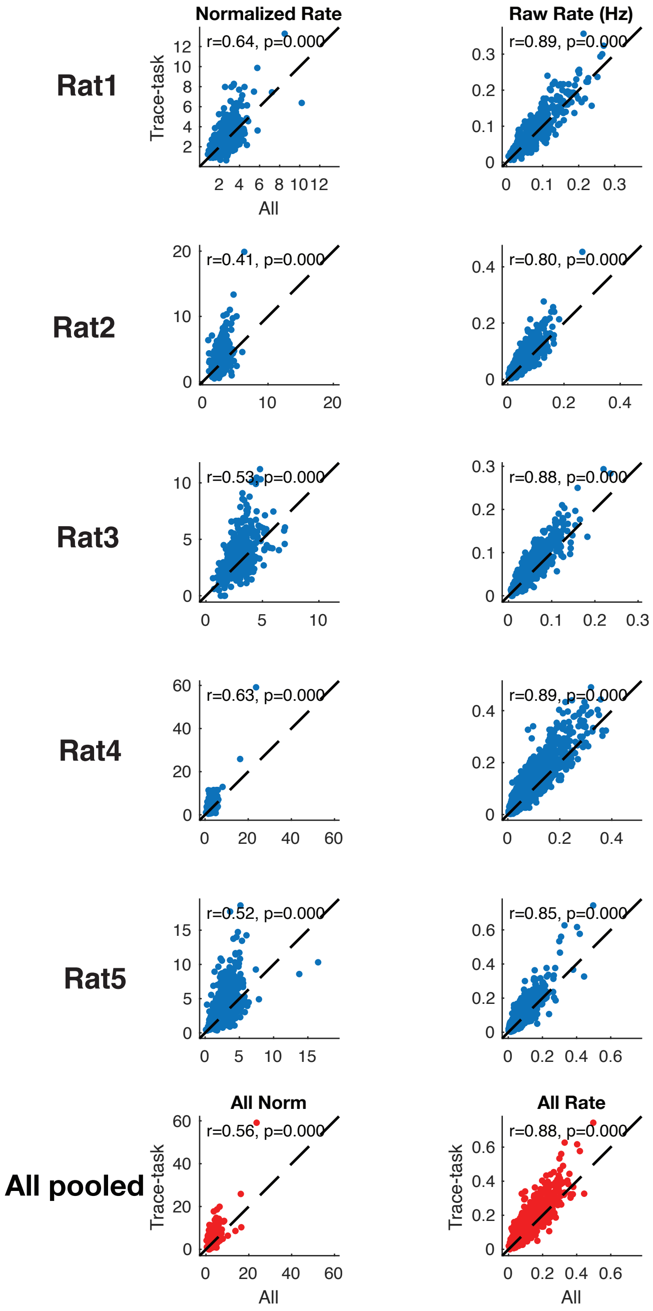

Normalized Rate
Raw Rate (Hz)
Rat1
Rat2
Rat3
Rat4
Rat5
All pooled
Trace-task
All
r=0.64, p=0.000
r=0.89, p=0.000
r=0.41, p=0.000
r=0.80, p=0.000
r=0.53, p=0.000
r=0.88, p=0.000
r=0.63, p=0.000
r=0.89, p=0.000
r=0.52, p=0.000
r=0.85, p=0.000
All Norm
All Rate
r=0.56, p=0.000
r=0.88, p=0.000

### Fig S5

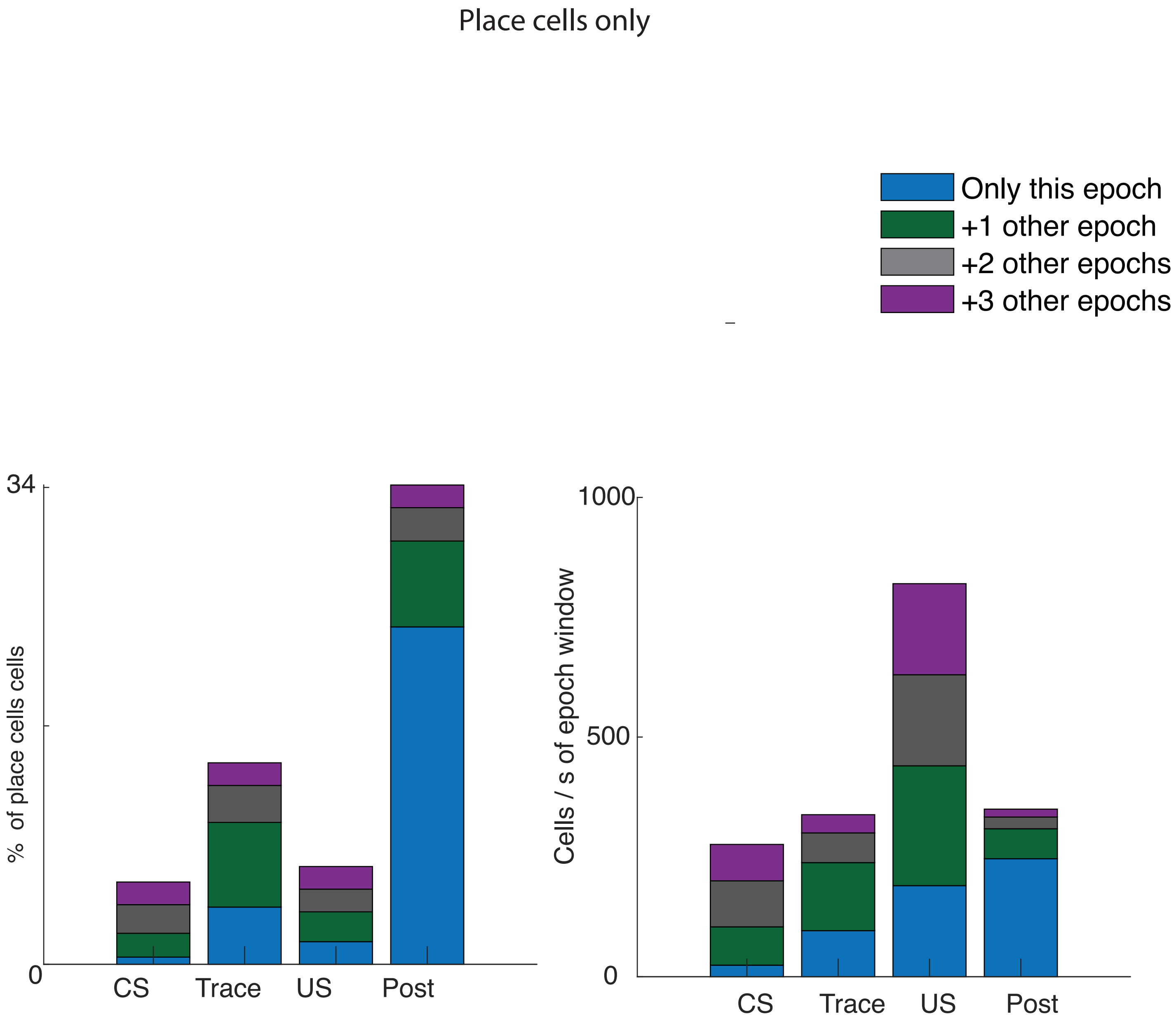
Place cells only
Only this epoch
+1 other epoch
+2 other epochs
+3 other epochs
34
% of place cells cells
0
CS
Trace
US
Post
1000
500
Cells / s of epoch window
0
CS
Trace
US
Post

### Fig S6

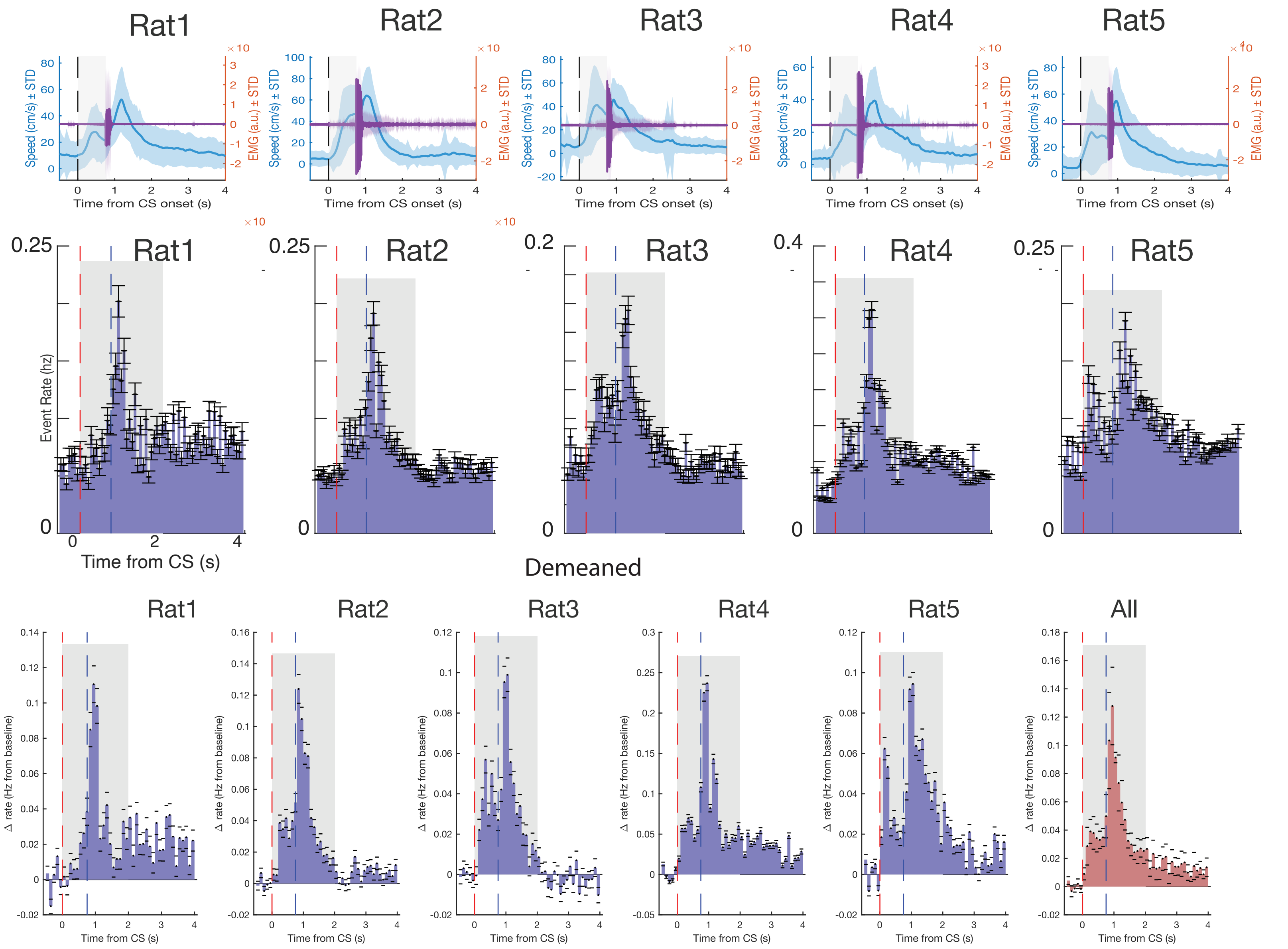

Rat1
Rat2
Rat3
Rat4
Rat5
Speed (cm/s) ± STD
EMG (a.u.) ± STD
Time from CS onset (s)
Event Rate (hz)
Time from CS (s)
Demeaned
All
Δ rate (Hz from baseline)

### Fig S7

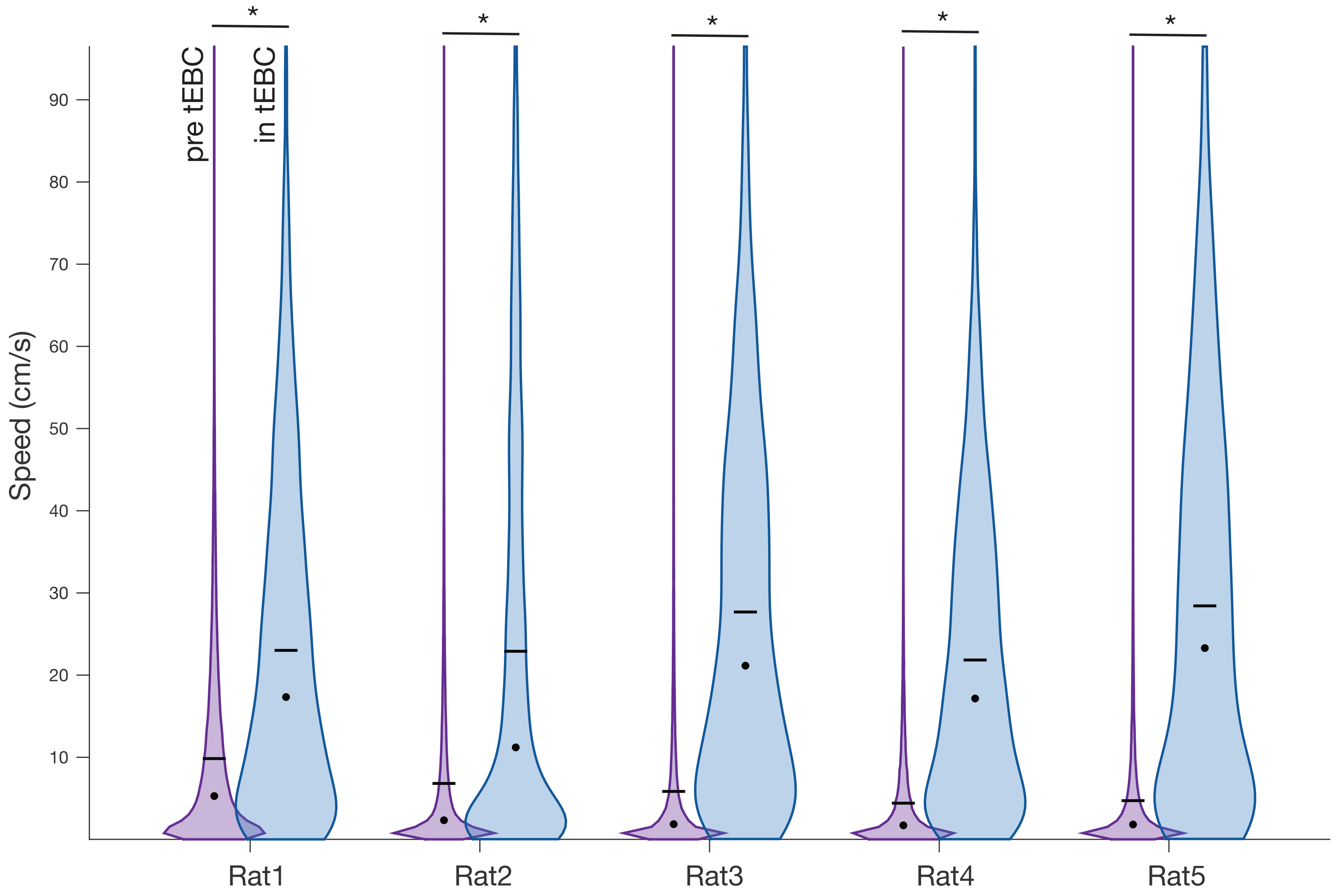

pre tEBC
in tEBC
*
*
*
*
*
Speed (cm/s)
90
80
70
60
50
40
30
20
10
Rat1
Rat2
Rat3
Rat4
Rat5

### Fig S8

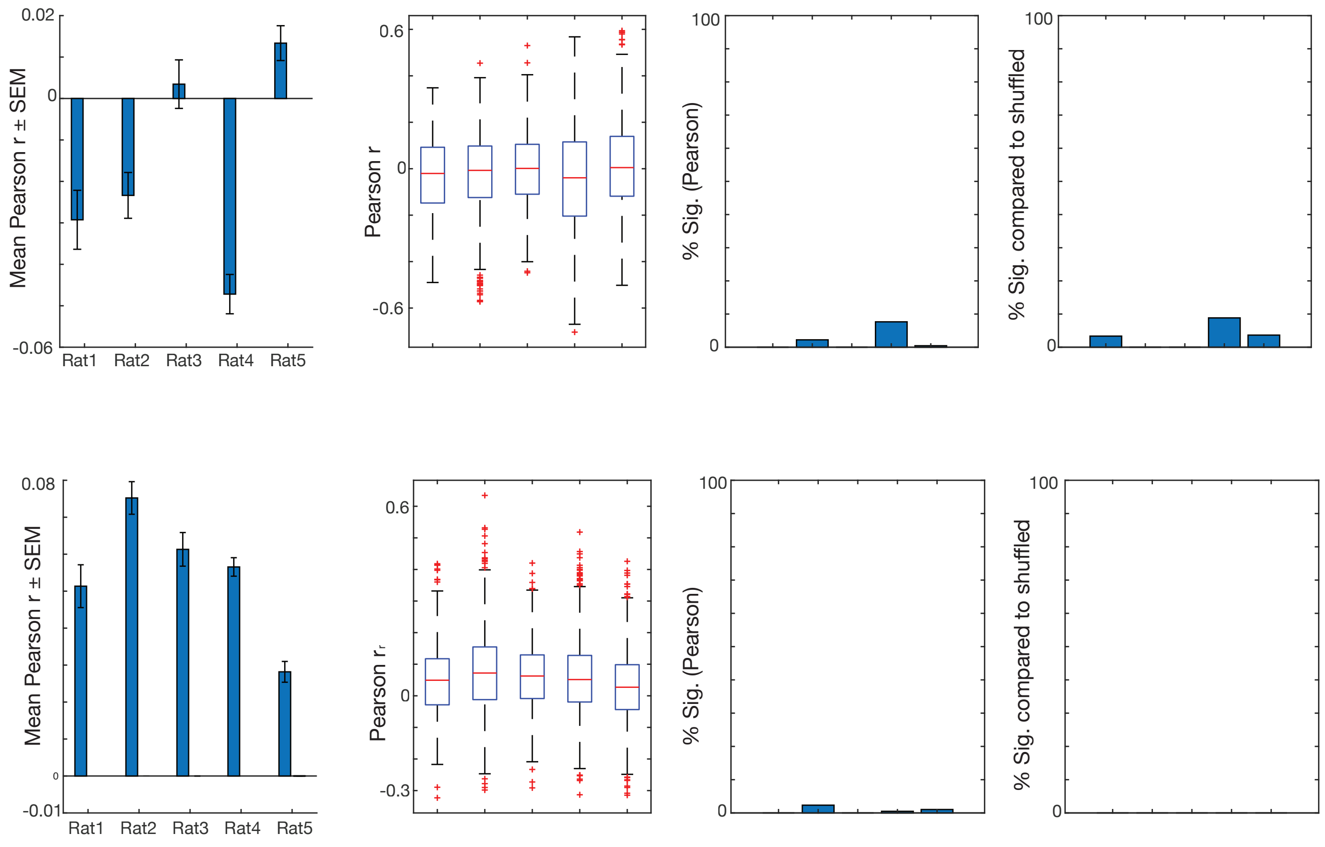

0.02
0
-0.06
Mean Pearson r ± SEM
Rat1
Rat2
Rat3
Rat4
Rat5
0.6
0
-0.6
Pearson r
100
0
% Sig. (Pearson)
100
0
% Sig. compared to shuffled
0.08
0
-0.01
Mean Pearson r ± SEM
Rat1
Rat2
Rat3
Rat4
Rat5
0.6
0
-0.3
Pearson r
100
0
% Sig. (Pearson)
100
0
% Sig. compared to shuffled

### Fig S9

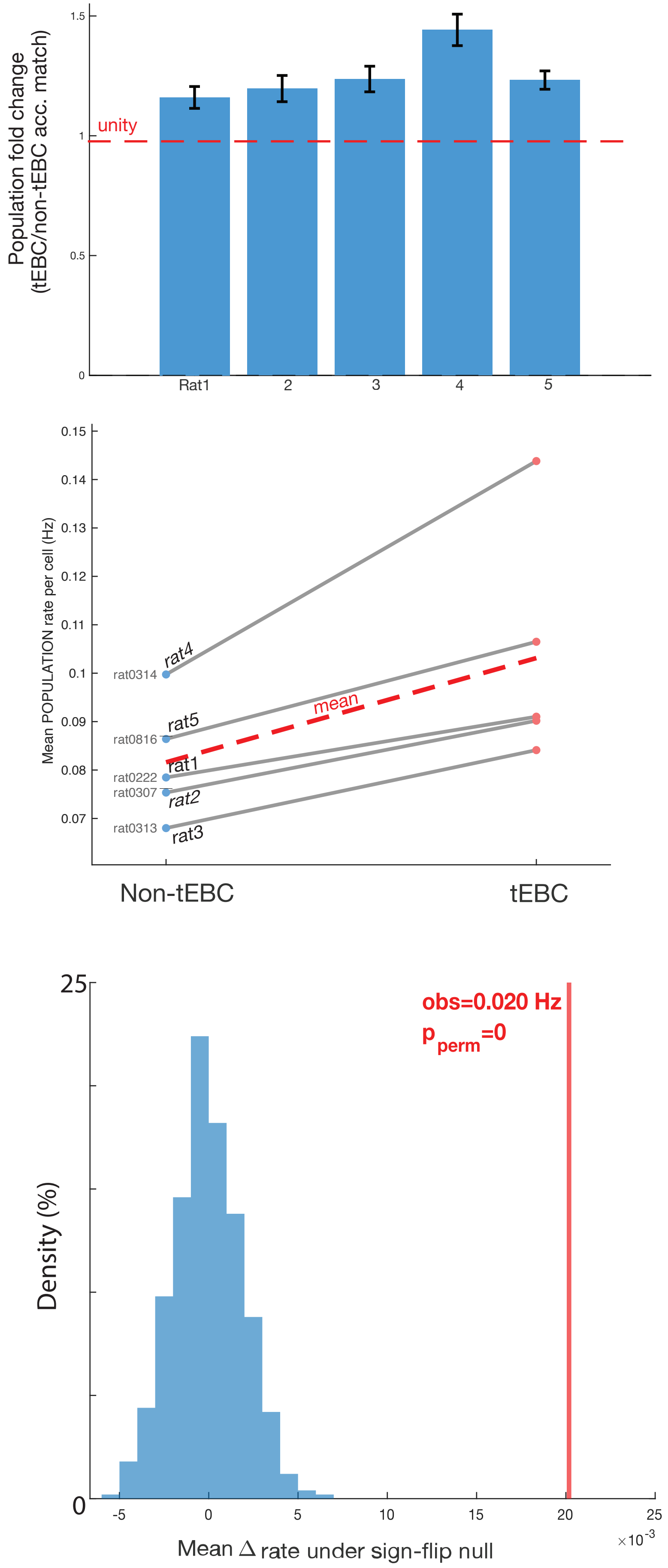

Population fold change
(tEBC/non-tEBC acc. match)
unity
1.5
1
0.5
0
Rat1
2
3
4
5
Mean POPULATION rate per cell (Hz)
0.15
0.14
0.13
0.12
0.11
0.1
0.09
0.08
0.07
rat0314
rat0816
rat0222
rat0307
rat0313
rat4
rat5
mean
rat1
rat2
rat3
Non-tEBC
tEBC
Density (%)
25
0
obs=0.020 Hz
$p_{perm}$=0
-5
0
5
10
15
20
25
$\times 10^{-3}$
Mean Δ rate under sign-flip null

### Fig S10

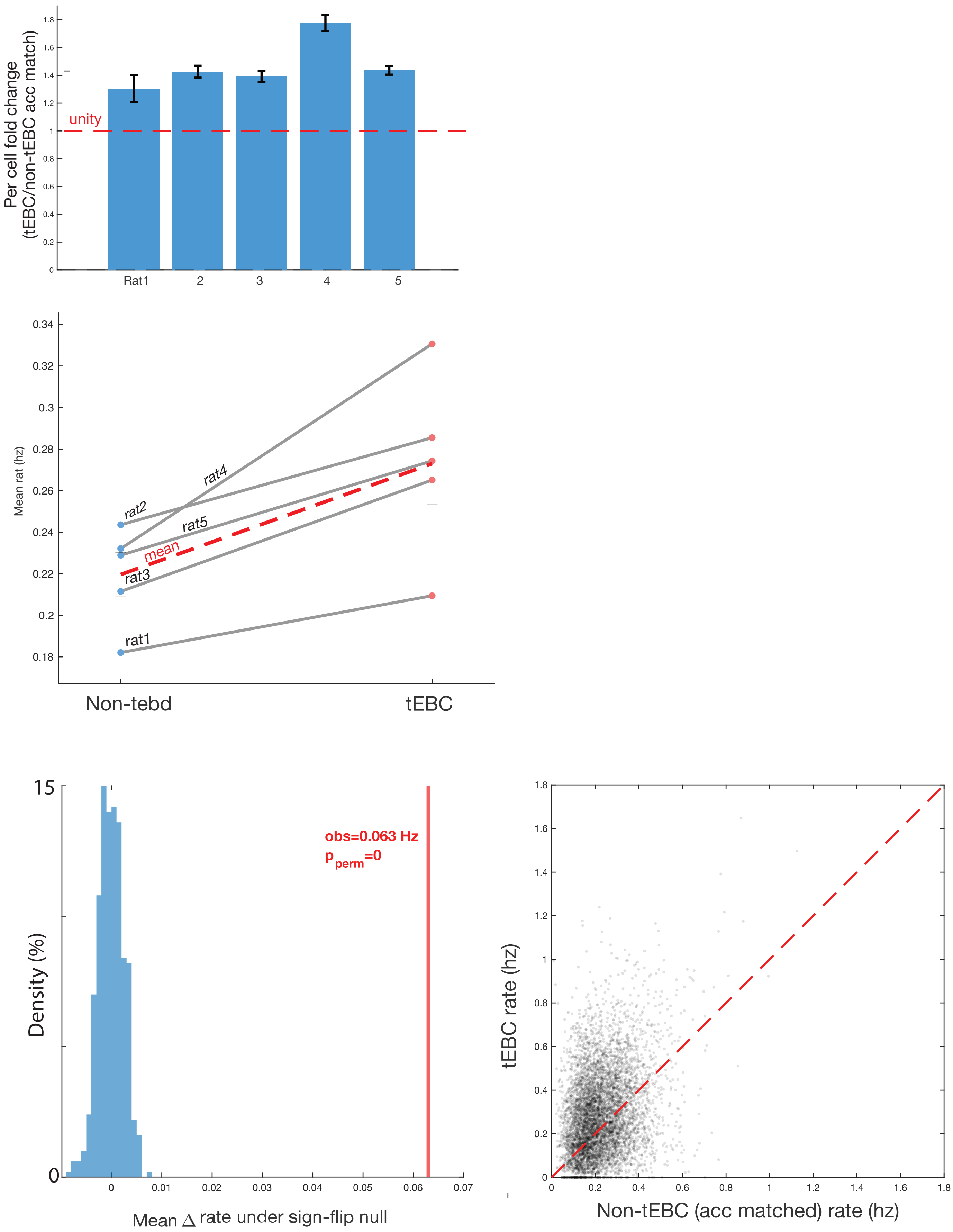
Per cell fold change (tEBC/non-tEBC acc match)
unity
Rat1
2
3
4
5
Mean rat (hz)
rat1
rat2
rat3
rat4
rat5
mean
Non-tebd
tEBC
Density (%)
obs=0.063 Hz
$p_{perm}$=0
Mean △ rate under sign-flip null
tEBC rate (hz)
Non-tEBC (acc matched) rate (hz)

### Fig S11

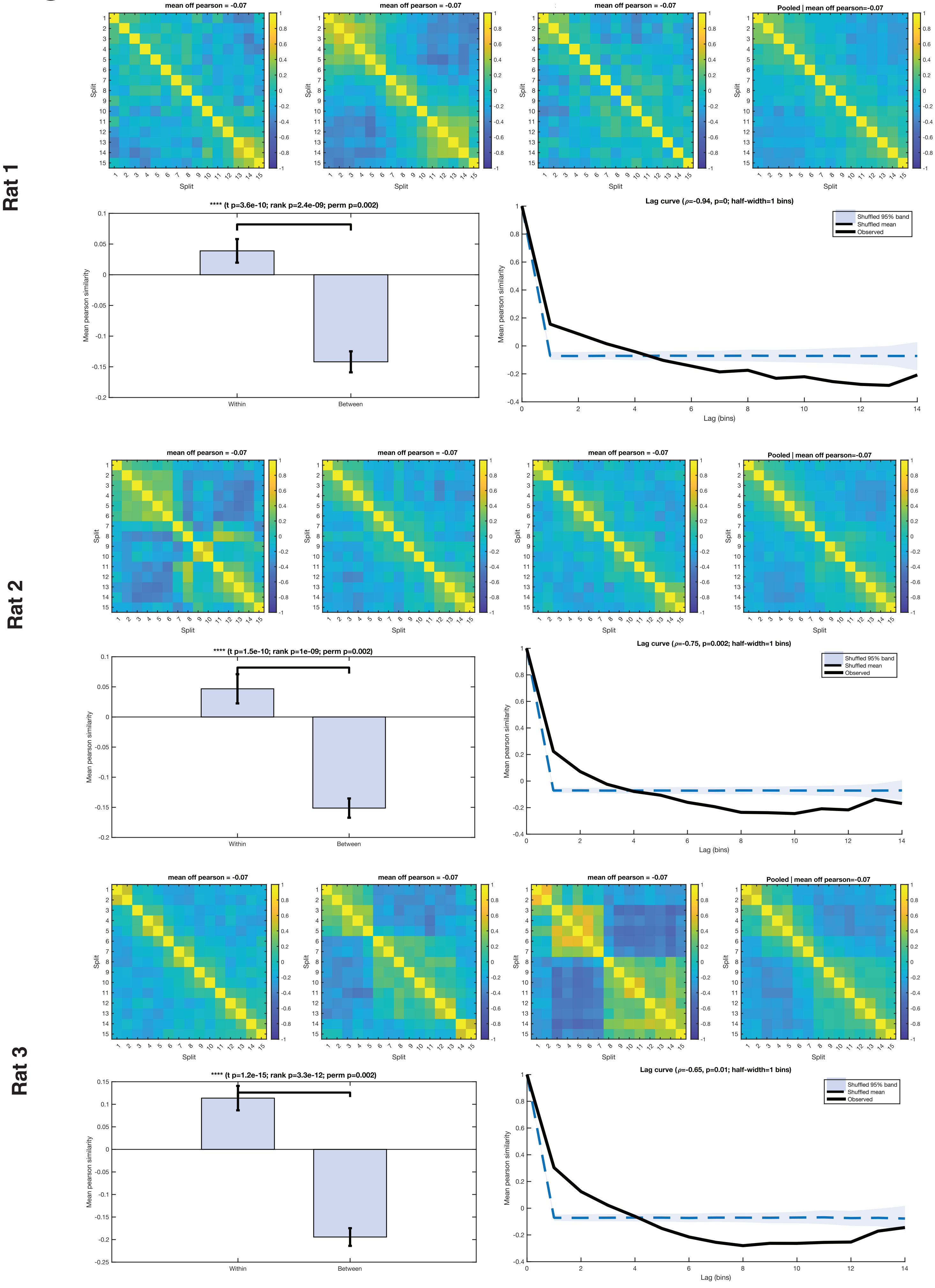
Rat 1
mean off pearson = -0.07
Pooled | mean off pearson=-0.07
Split
**** (t p=3.6e-10; rank p=2.4e-09; perm p=0.002)
Mean pearson similarity
Within
Between
Lag curve (ρ=-0.94, p=0; half-width=1 bins)
Shuffled 95% band
Shuffled mean
Observed
Lag (bins)
Rat 2
**** (t p=1.5e-10; rank p=1e-09; perm p=0.002)
Lag curve (ρ=-0.75, p=0.002; half-width=1 bins)
Rat 3
**** (t p=1.2e-15; rank p=3.3e-12; perm p=0.002)
Lag curve (ρ=-0.65, p=0.01; half-width=1 bins)

### Fig S11

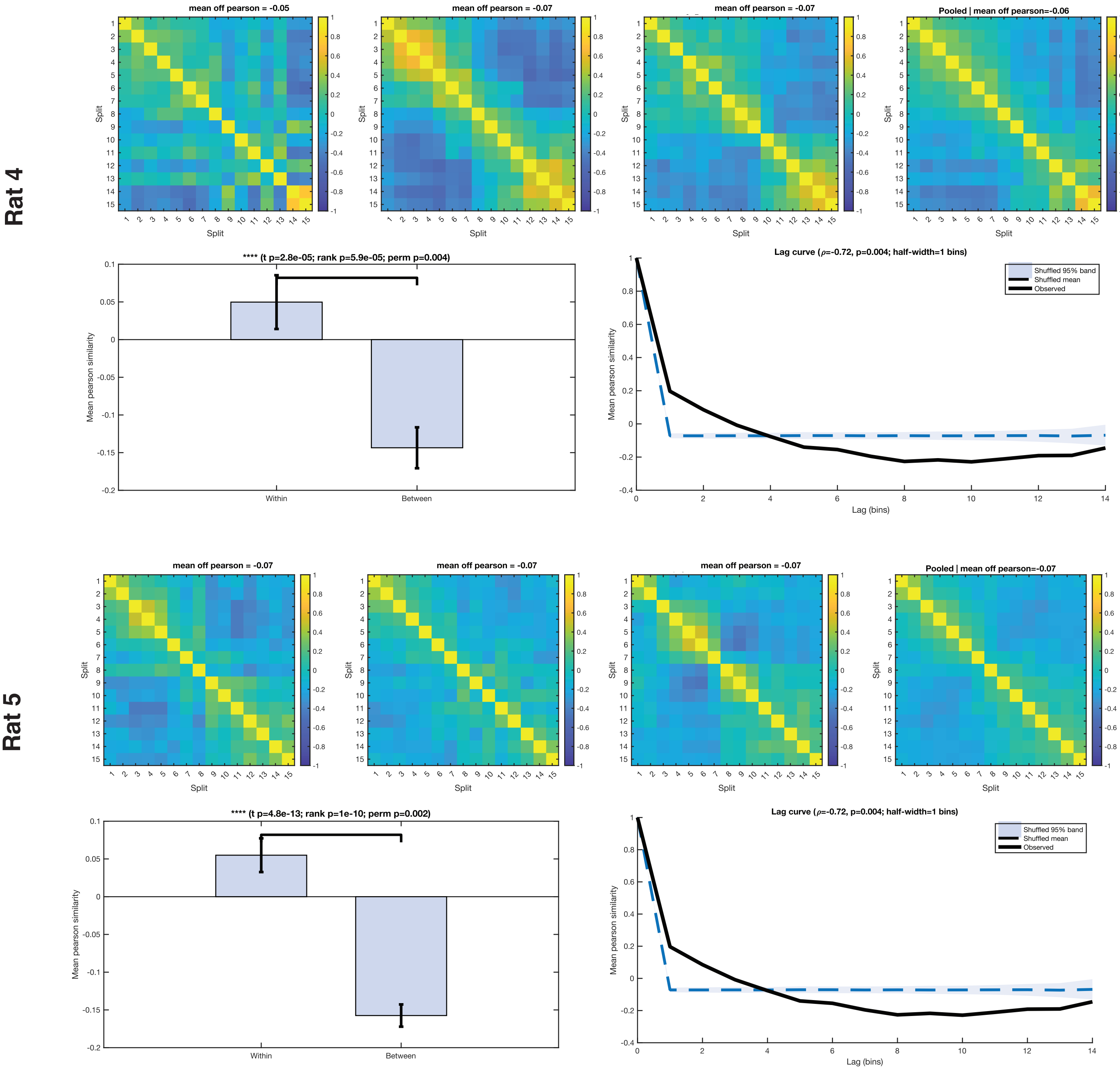

Rat 4
mean off pearson = -0.05
mean off pearson = -0.07
mean off pearson = -0.07
Pooled | mean off pearson=-0.06
Split
**** (t p=2.8e-05; rank p=5.9e-05; perm p=0.004)
Mean pearson similarity
Within
Between
Lag curve (ρ=-0.72, p=0.004; half-width=1 bins)
Shuffled 95% band
Shuffled mean
Observed
Lag (bins)
Rat 5
mean off pearson = -0.07
mean off pearson = -0.07
mean off pearson = -0.07
Pooled | mean off pearson=-0.07
**** (t p=4.8e-13; rank p=1e-10; perm p=0.002)
Lag curve (ρ=-0.72, p=0.004; half-width=1 bins)

### Fig S12

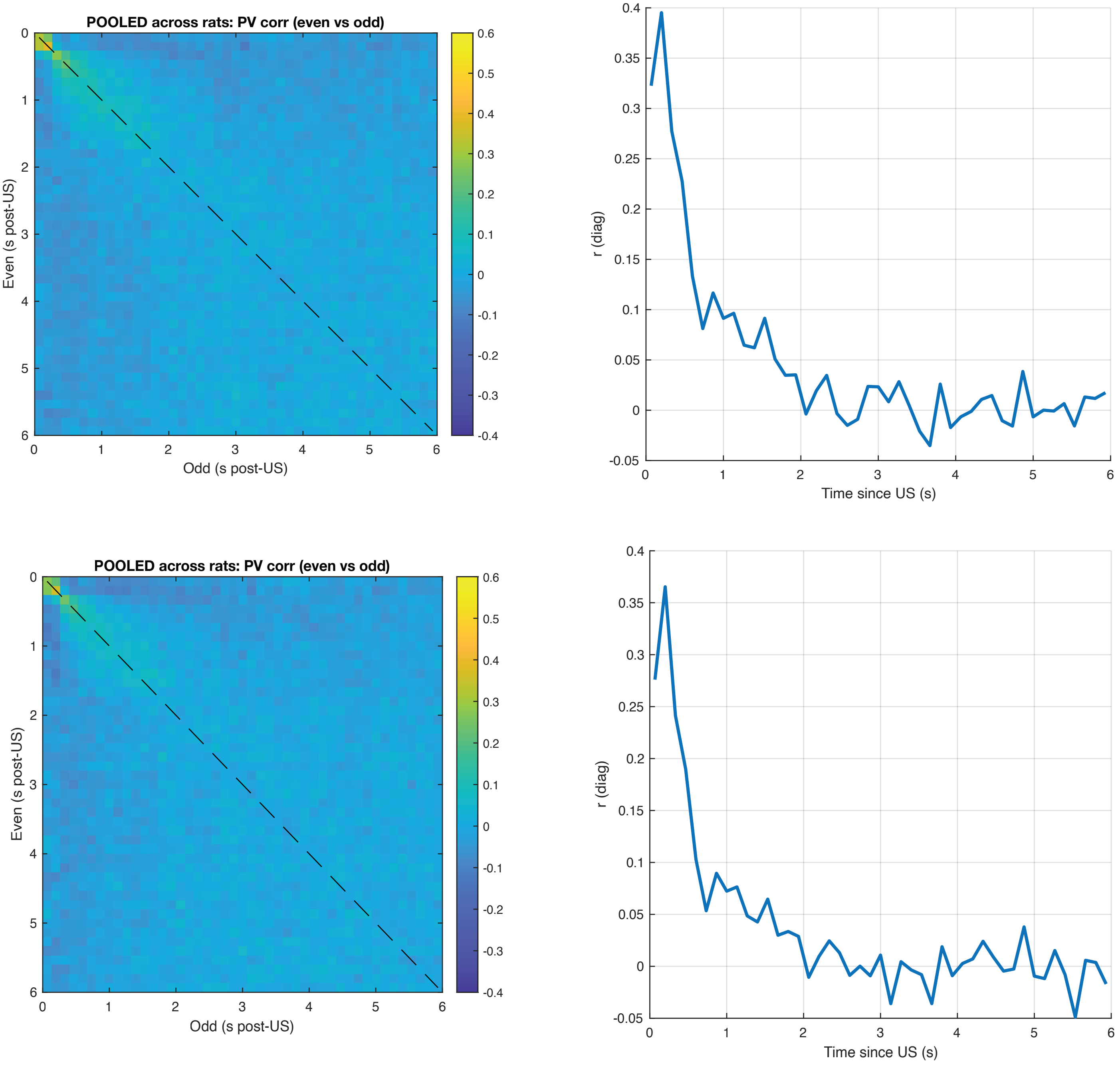

POOLED across rats: PV corr (even vs odd)
Even (s post-US)
Odd (s post-US)
r (diag)
Time since US (s)
POOLED across rats: PV corr (even vs odd)
Even (s post-US)
Odd (s post-US)
r (diag)
Time since US (s)

### Fig S13

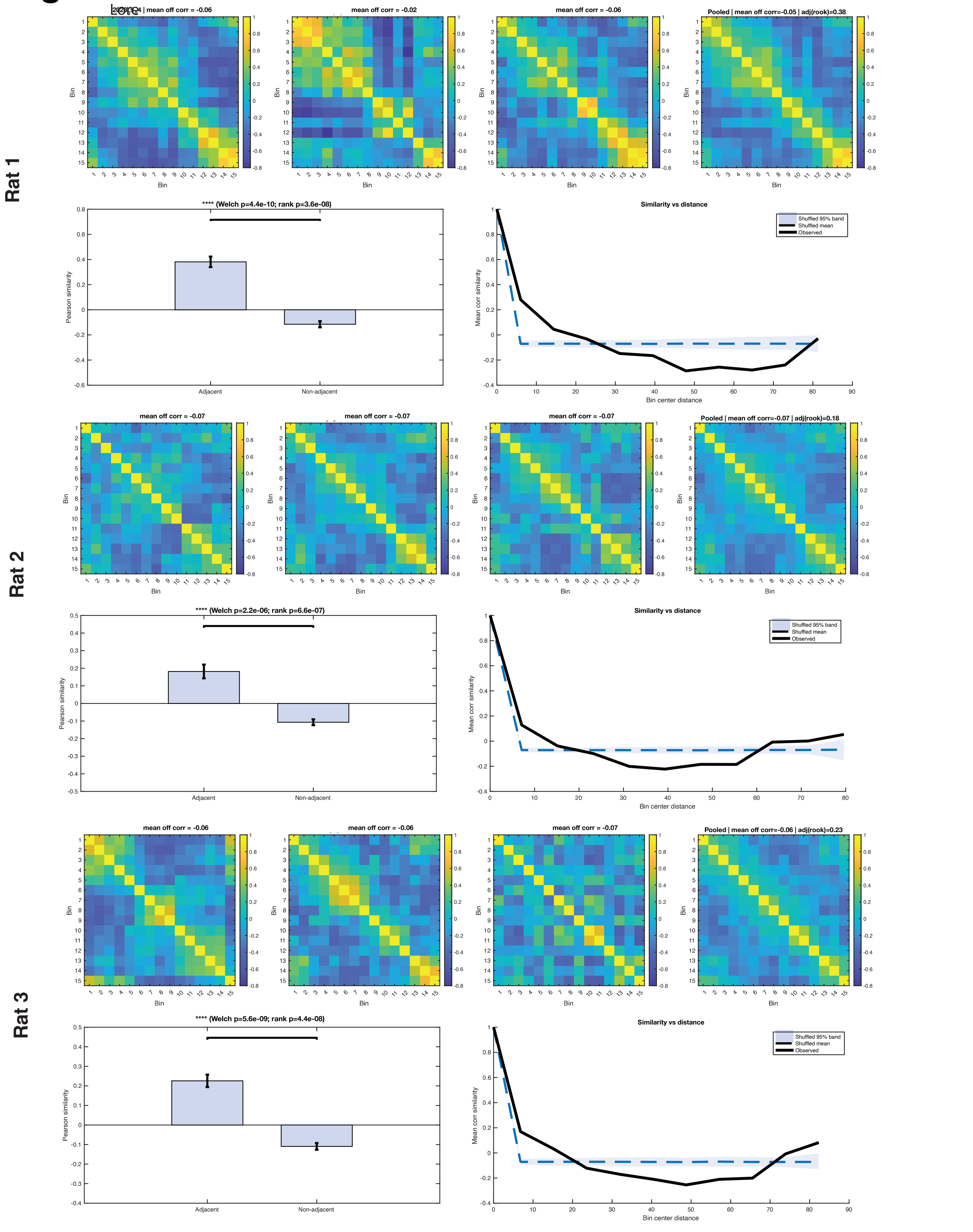

Rat 1
mean off corr = -0.06
mean off corr = -0.02
mean off corr = -0.06
Pooled | mean off corr=-0.05 | adj(rook)=0.38
Bin
**** (Welch p=4.4e-10; rank p=3.6e-08)
Pearson similarity
Adjacent
Non-adjacent
Similarity vs distance
Shuffled 95% band
Shuffled mean
Observed
Mean corr similarity
Bin center distance
Rat 2
mean off corr = -0.07
mean off corr = -0.07
mean off corr = -0.07
Pooled | mean off corr=-0.07 | adj(rook)=0.18
**** (Welch p=2.2e-06; rank p=6.6e-07)
Rat 3
mean off corr = -0.06
mean off corr = -0.06
mean off corr = -0.07
Pooled | mean off corr=-0.06 | adj(rook)=0.23
**** (Welch p=5.6e-09; rank p=4.4e-08)

### Fig S13

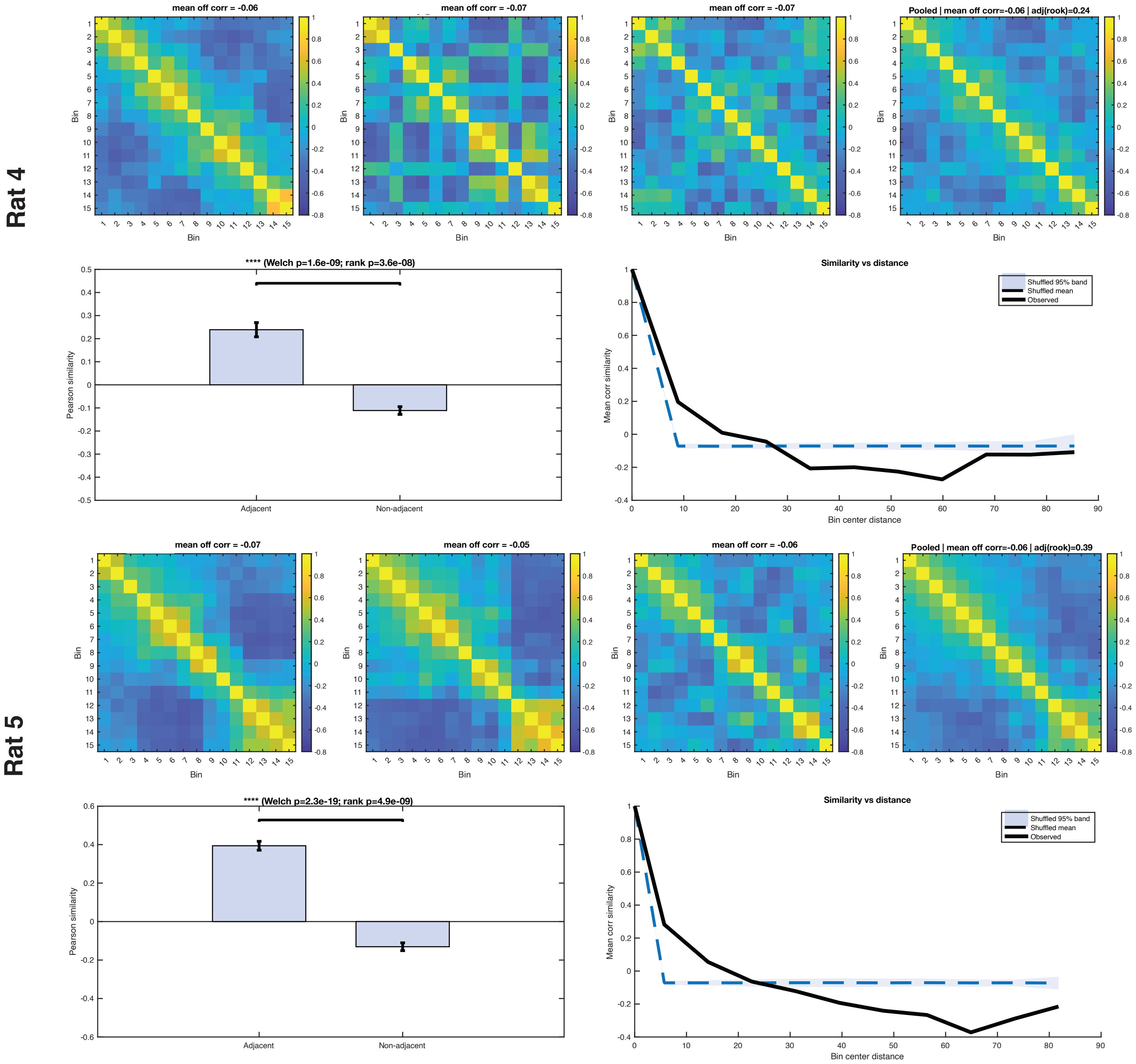

Rat 4
mean off corr = -0.06
mean off corr = -0.07
mean off corr = -0.07
Pooled | mean off corr=-0.06 | adj(rook)=0.24
Bin
**** (Welch p=1.6e-09; rank p=3.6e-08)
Pearson similarity
Adjacent
Non-adjacent
Similarity vs distance
Shuffled 95% band
Shuffled mean
Observed
Mean corr similarity
Bin center distance
Rat 5
mean off corr = -0.07
mean off corr = -0.05
mean off corr = -0.06
Pooled | mean off corr=-0.06 | adj(rook)=0.39
Bin
**** (Welch p=2.3e-19; rank p=4.9e-09)
Pearson similarity
Adjacent
Non-adjacent
Similarity vs distance
Shuffled 95% band
Shuffled mean
Observed
Mean corr similarity
Bin center distance

### Fig S14

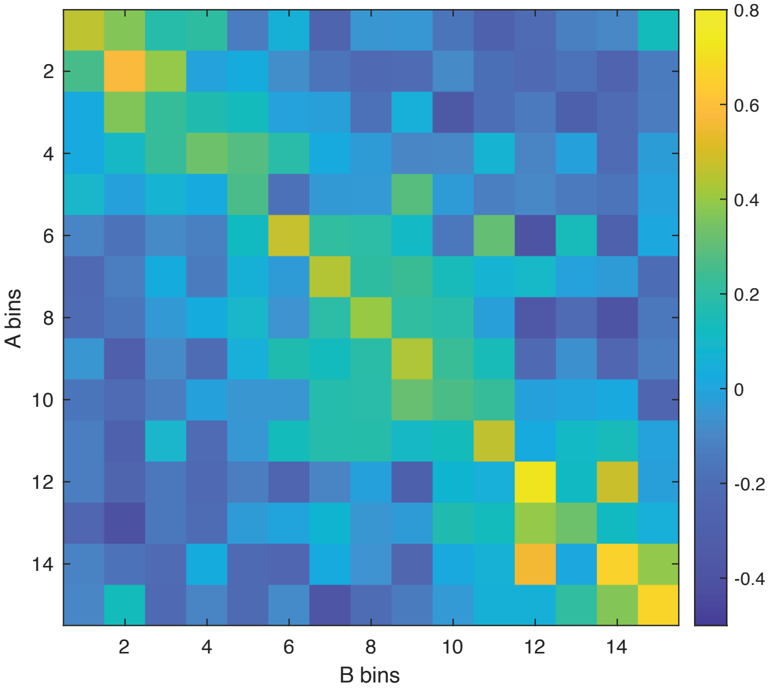

A bins
B bins
2
4
6
8
10
12
14
0.8
0.6
0.4
0.2
0
-0.2
-0.4


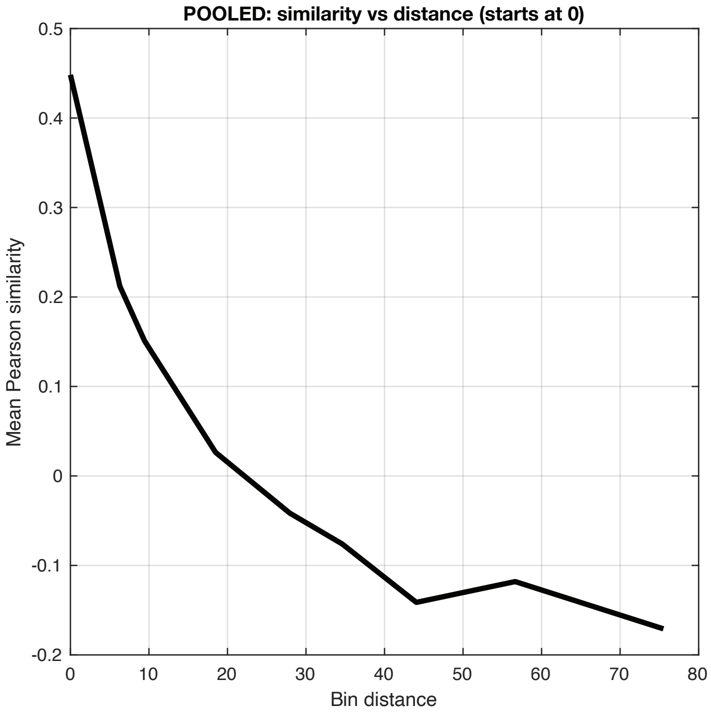

POOLED: similarity vs distance (starts at 0)
Mean Pearson similarity
Bin distance
0.5
0.4
0.3
0.2
0.1
0
-0.1
-0.2
0
10
20
30
40
50
60
70
80

### Fig S15

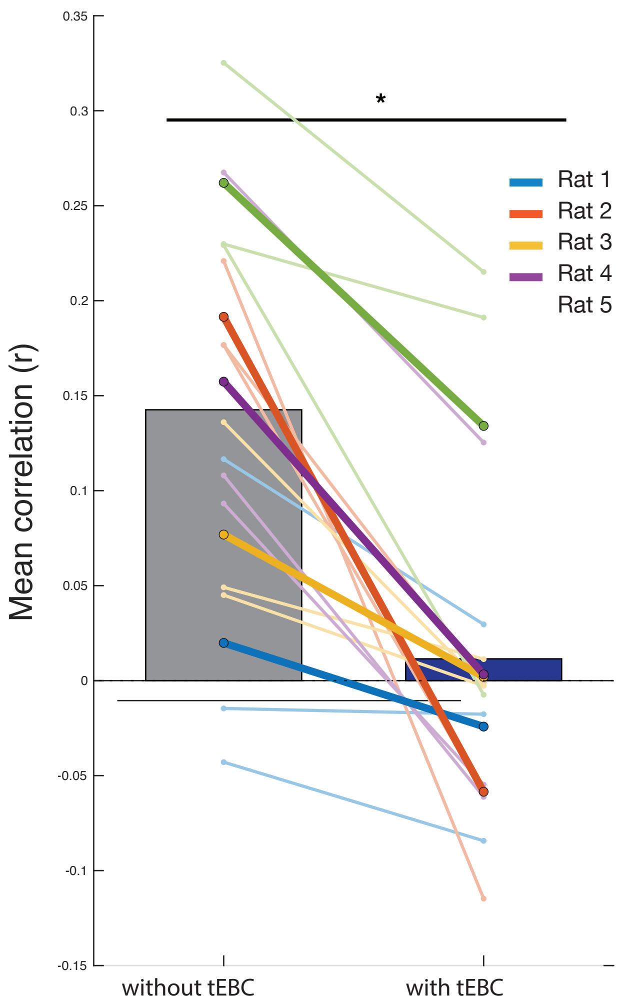

0.35
0.3
0.25
0.2
0.15
0.1
0.05
0
-0.05
-0.1
-0.15
Mean correlation (r)
*
Rat 1
Rat 2
Rat 3
Rat 4
Rat 5
without tEBC
with tEBC

### Fig S16

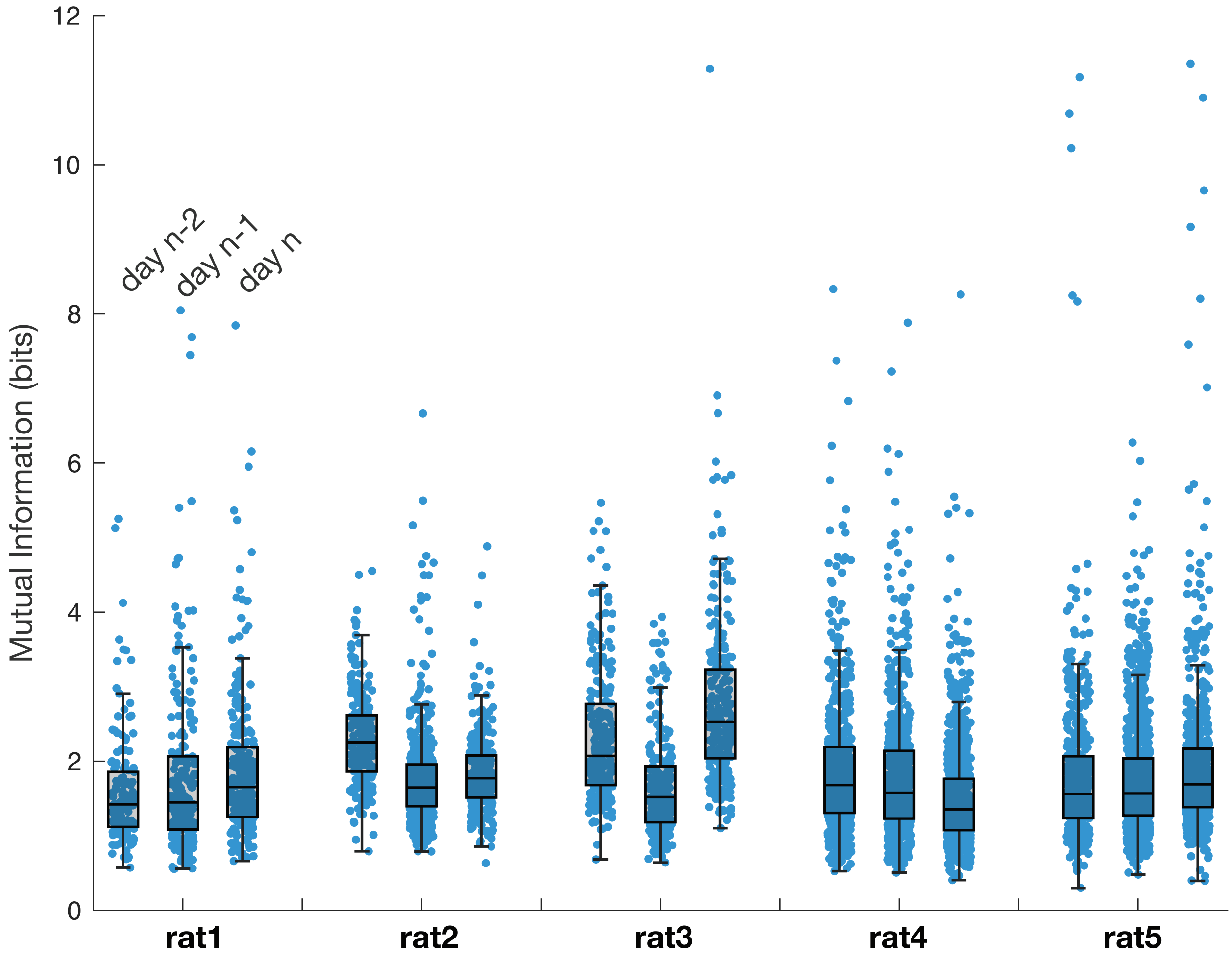
Mutual Information (bits)
0
2
4
6
8
10
12
day n-2
day n-1
day n
rat1
rat2
rat3
rat4
rat5

### Fig S17

## a

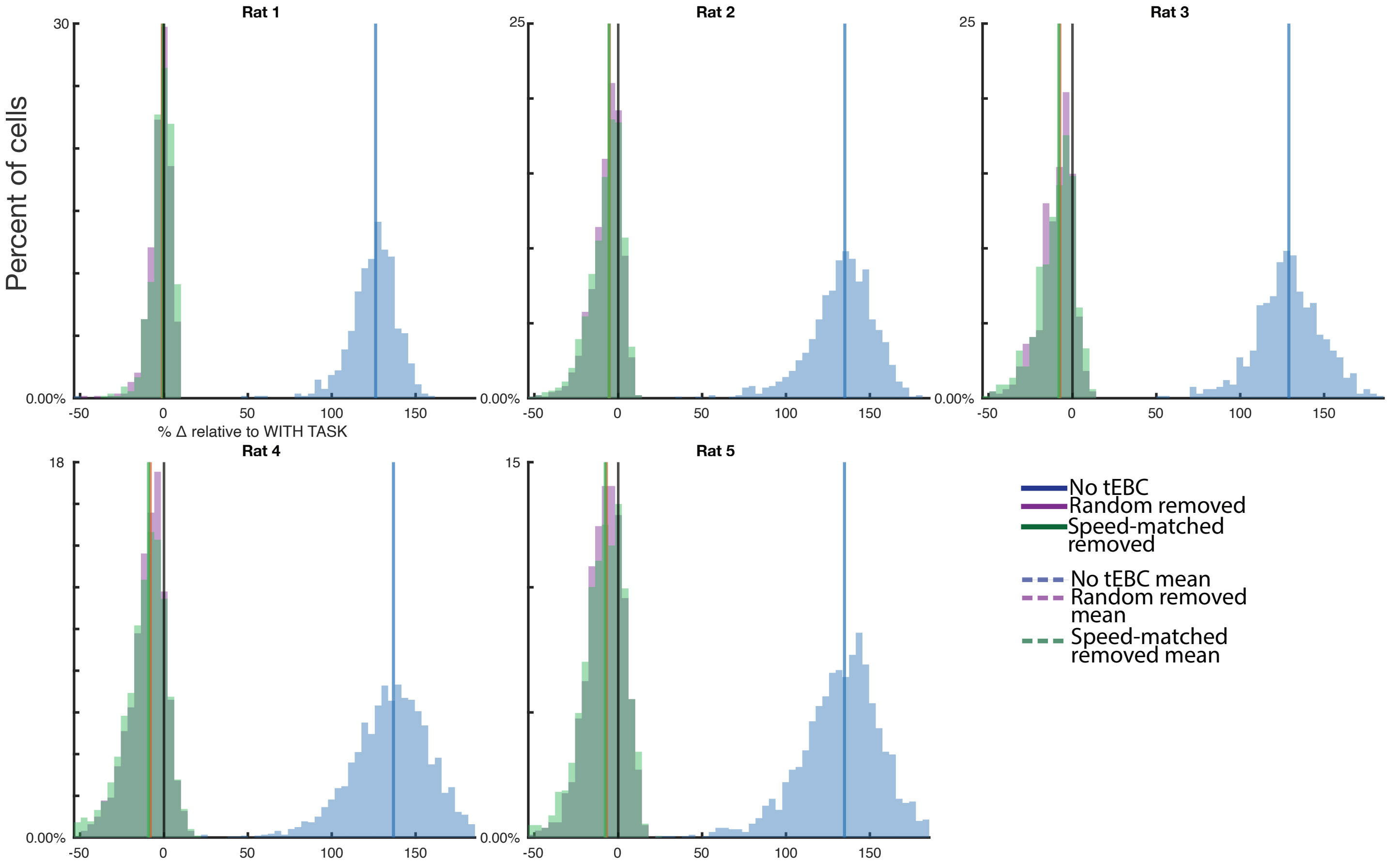

Rat 1
Rat 2
Rat 3
Rat 4
Rat 5
Percent of cells
% Δ relative to WITH TASK
30
25
25
18
15
0.00%
-50
0
50
100
150
No tEBC
Random removed
Speed-matched removed
No tEBC mean
Random removed mean
Speed-matched removed mean

### Fig S18

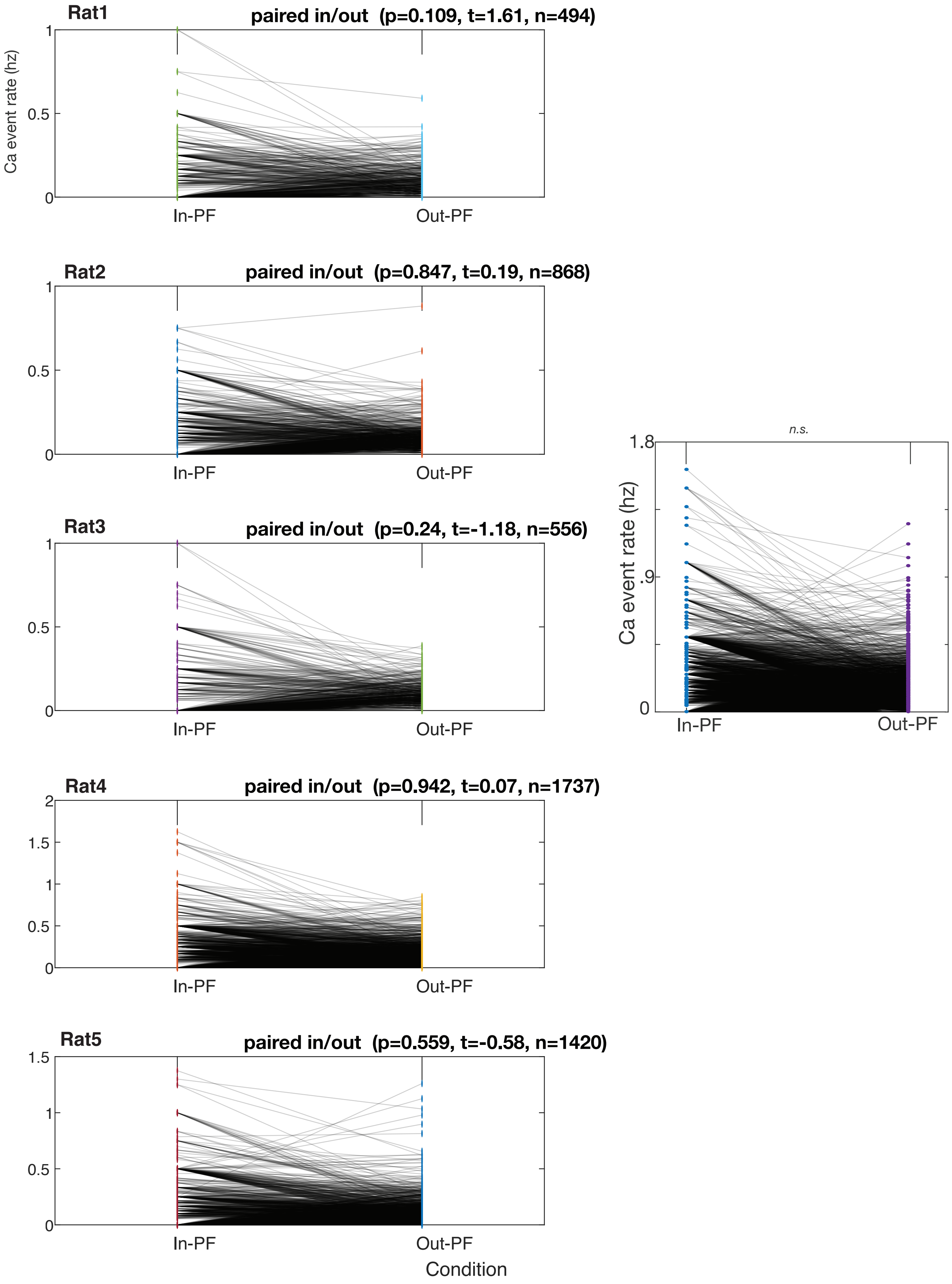
Rat1
paired in/out (p=0.109, t=1.61, n=494)
Ca event rate (hz)
1
0.5
0
In-PF
Out-PF
Rat2
paired in/out (p=0.847, t=0.19, n=868)
1
0.5
0
In-PF
Out-PF
Rat3
paired in/out (p=0.24, t=-1.18, n=556)
1
0.5
0
In-PF
Out-PF
Rat4
paired in/out (p=0.942, t=0.07, n=1737)
2
1.5
1
0.5
0
In-PF
Out-PF
Rat5
paired in/out (p=0.559, t=-0.58, n=1420)
1.5
1
0.5
0
In-PF
Out-PF
Condition
n.s.
1.8
.9
0
Ca event rate (hz)
In-PF
Out-PF

### Fig S19

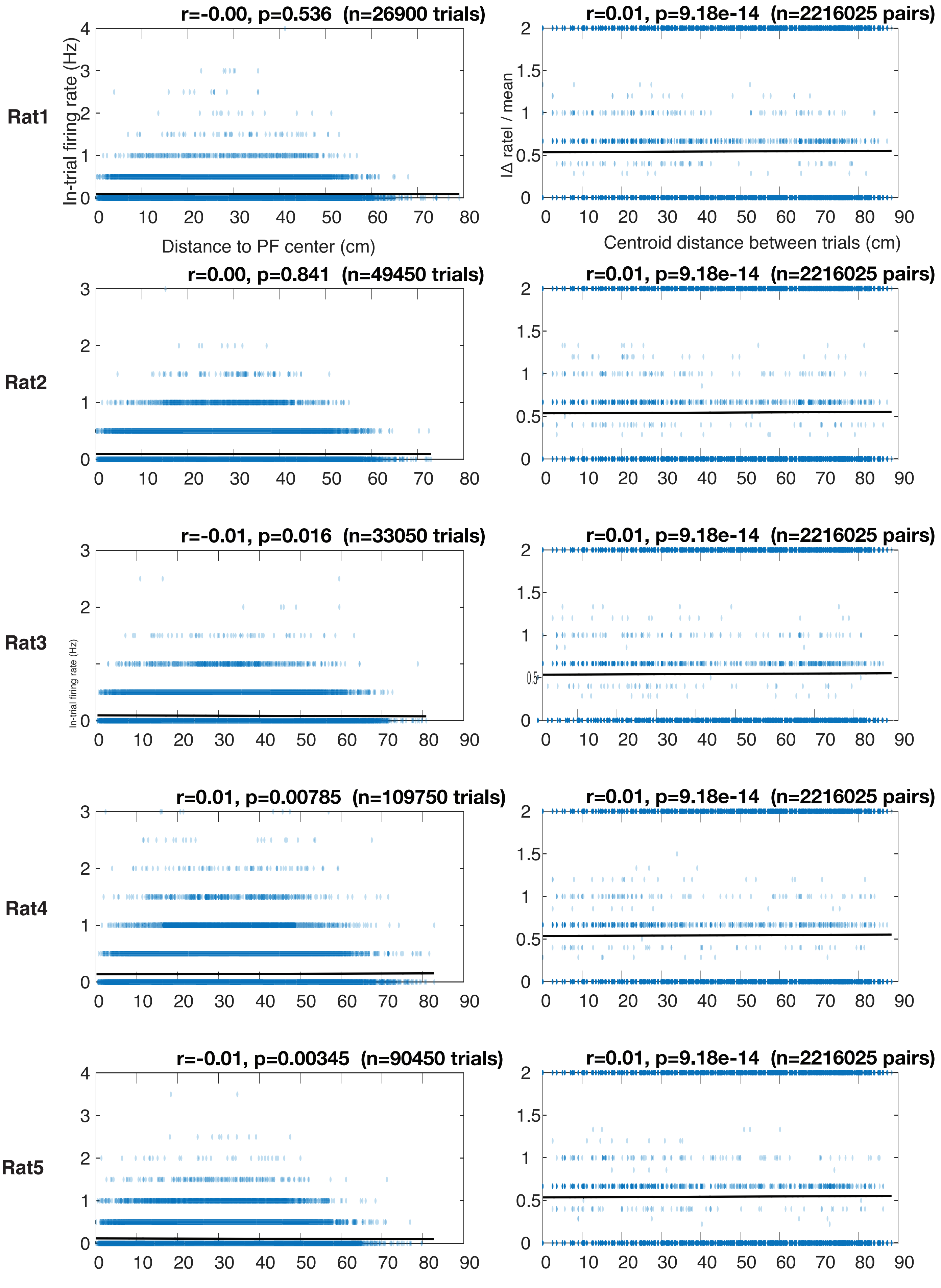

Rat1
r=-0.00, p=0.536 (n=26900 trials)
In-trial firing rate (Hz)
Distance to PF center (cm)
r=0.01, p=9.18e-14 (n=2216025 pairs)
|Δ rate| / mean
Centroid distance between trials (cm)
Rat2
r=0.00, p=0.841 (n=49450 trials)
r=0.01, p=9.18e-14 (n=2216025 pairs)
Rat3
r=-0.01, p=0.016 (n=33050 trials)
In-trial firing rate (Hz)
r=0.01, p=9.18e-14 (n=2216025 pairs)
Rat4
r=0.01, p=0.00785 (n=109750 trials)
r=0.01, p=9.18e-14 (n=2216025 pairs)
Rat5
r=-0.01, p=0.00345 (n=90450 trials)
r=0.01, p=9.18e-14 (n=2216025 pairs)

### Fig S20

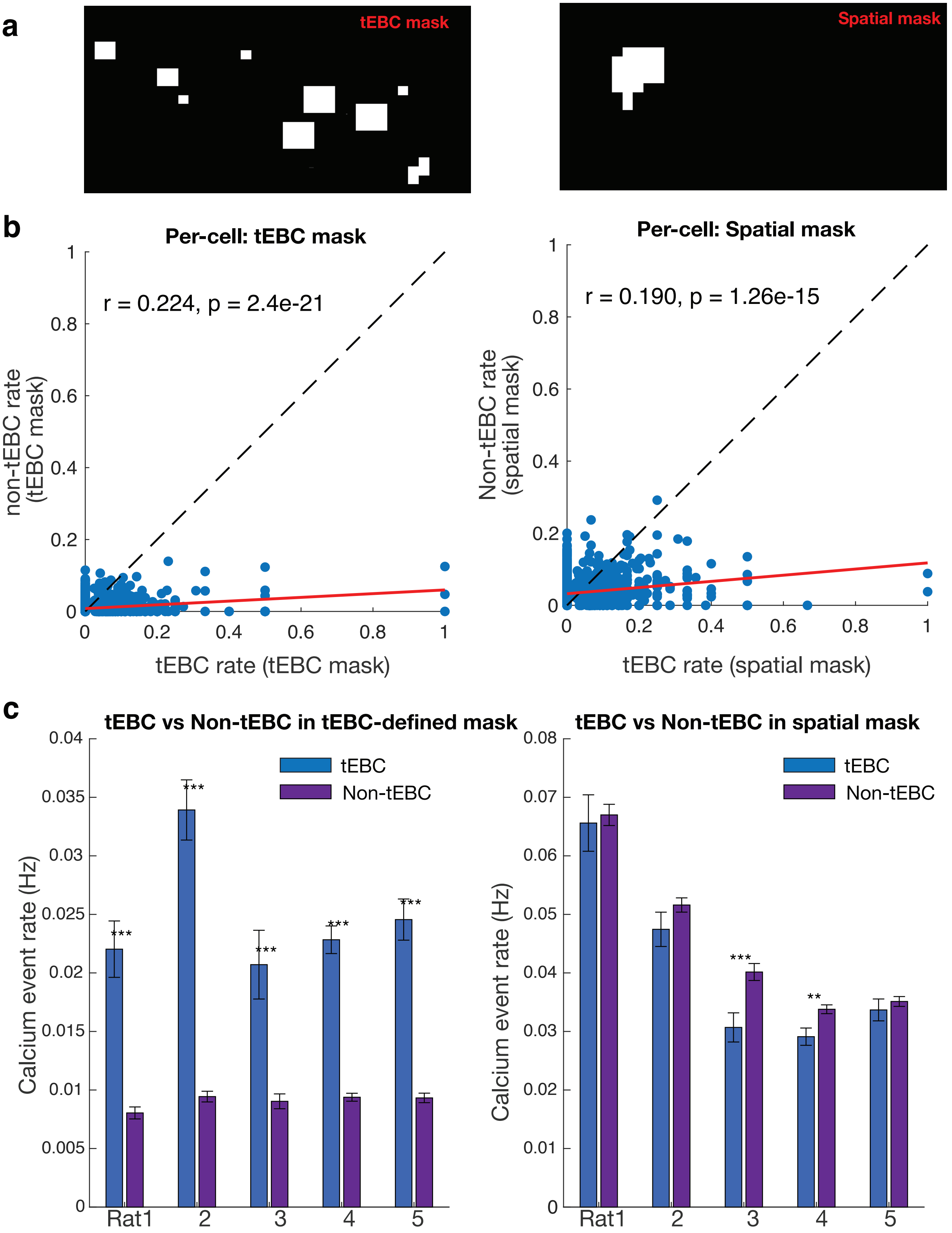
a
tEBC mask
Spatial mask
b
Per-cell: tEBC mask
r = 0.224, p = 2.4e-21
non-tEBC rate (tEBC mask)
tEBC rate (tEBC mask)
Per-cell: Spatial mask
r = 0.190, p = 1.26e-15
Non-tEBC rate (spatial mask)
tEBC rate (spatial mask)
c
tEBC vs Non-tEBC in tEBC-defined mask
tEBC
Non-tEBC
Calcium event rate (Hz)
Rat1
2
3
4
5
tEBC vs Non-tEBC in spatial mask
tEBC
Non-tEBC
Calcium event rate (Hz)
Rat1
2
3
4
5
